# Reversing Transcriptome-Wide Association Studies to improve expression Quantitative Trait Loci associations

**DOI:** 10.1101/2020.12.27.424460

**Authors:** Jie Yuan, Ben Lai, Itsik Pe’er

## Abstract

Transcriptome-Wide Association Studies discover SNP effects mediated by gene expression through a two-stage process: a typically small reference panel is used to infer SNP-expression effects, and then these are applied to discover associations between imputed expression and phenotypes. We investigate whether the accuracy of SNP-expression and expression-phenotype associations can be increased by performing inference on both the reference panel and independent GWAS cohorts simultaneously. We develop EMBER (Estimation of Mediated Binary Effects in Regression) to re-estimate these effects using a liability threshold model with an adjustment to variance components accounting for imputed expression from GWAS data. In simulated data with only gene-mediated effects, EMBER more than doubles the performance of SNP-expression linear regression, increasing mean *r*^2^ from 0.3 to 0.65 with a gene-mediated variance explained of 0.01. EMBER also improves estimation accuracy when the fraction of cis-SNP variance mediated by genes is as low as 30%. We apply EMBER to genotype and gene expression data in schizophrenia by combining 512 samples from the CommonMind Consortium and 56,081 samples from the Psychiatric Genomic Consortium. We evaluate performance of EMBER in 36 genes suggested by TWAS by concordance of inferred effects with effects reported independently for frontal cortex expression. Applying the EMBER framework to a baseline linear regression model increases performance in 26 out of 36 genes (sign test p-value .0020) with an increase in mean *r*^2^ from 0.200 to 0.235.

## 2 Introduction

Genome-wide association methods have to date discovered thousands of associations with disease risks and other phenotypes [1]. However, functional studies to determine the causal genes or mechanisms behind these associations have yet to be fully explored [2]. One such strategy is to identify expression Quantitative Trait Loci (eQTLs), variants in the genome which are implicated in mRNA expression. eQTLs have been shown to be enriched in GWAS-significant SNPs and further enrichment is found over fine-mapped cis-eQTLs both across all tissues and for specific pairs of tissues and phenotypes [3, 4]. Large consortia such as GTEx [5, 6] have discovered numerous eQTLs by repeating single-variant regressions over cis-SNPs [7, 8], though challenges remain such as low power and winner’s curse stemming from small sample sizes of current RNA-seq experiments [9].

Transcriptome-Wide Association Studies (TWAS) have played a promising role in discovering eQTL associations with disease risk [10, 11]. The method uses reference panels of known SNP-gene effect sizes obtained from smaller gene expression experiments to predict associations between expression and disease phenotypes. SNP-gene effects are trained using multi-SNP regression methods: Gusev et al. [11] use primarily the best linear unbiased predictor (BLUP) whereas the PrediXcan model by Gamazon et al. [10] uses elastic net. Associations between imputed gene expression and disease phenotypes are then assessed either by imputing expression for each individual in large GWAS cohorts or computed directly from summary statistic information. While TWAS performs only a Z-score to test imputed gene-trait association, other iterations of this framework include MR-Egger, a method for estimating of the imputed gene effect size through Mendelian randomization, which accounts for direct SNP-trait effects not mediated by genes, and further refinements of these methods to factor out correlations between SNPs due to linkage disequilibrium [12]. To date, TWAS has discovered a large number of imputed gene-phenotype associations in a wide variety of phenotypes including schizophrenia [13]. However, challenges to interpretation remain, as TWAS associations are not necessarily causal, and genes may be mistakenly tagged due to pleiotropy at the SNP level. Additional noise is provided by small sample sizes in SNP-expression reference panels [14].

We developed EMBER (Estimation of Mediated Binary Effects in Regression) to investigate whether SNP-gene and gene-trait effect sizes in TWAS can be improved by performing inference on both a reference panel and GWAS data simultaneously, rather than a two-step process in which eQTLs are first inferred, then assumed to be fixed and true. As this model must account for both observed gene expression from the reference panel and imputed gene expression from an estimate of eQTLs, EMBER applies the liability threshold model and its treatment of variance explained to represent differing levels of confidence in gene expression estimates [15]. The liability threshold model assumes an observed binary trait, such as disease status, is sampled by thresholding a latent normally distributed liability function. The variance explained by a set of predictors, such as SNPs or imputed gene expression, is then a fraction of the variance of the standard normal liability distribution.

We find in simulations that EMBER significantly improves estimation of SNP-expression effects when SNP effects are mediated by gene expression. Even when non-mediated effects are present, EMBER can still improve SNP-expression estimation when the variance explained of mediated effects is at least 30%. In practice, we obtained the largest boost in SNP-expression estimation performance from a variable reduction preprocessing step, far outperforming the BLUP and elastic net on schizoprenia cohorts from the Common-Mind Consortium [16] and the Schizophrenia Working Group of the Psychiatric Genomics Consortium [17]. After the variable reduction step, we evaluate EMBER in 36 genes with significant TWAS effects as reported by Gusev et al. [11]. While most of these genes mediate far less than 30% of the variance explained of their cis-SNPs, we nevertheless find that for a subset of these genes, EMBER results in slight improvements in *r*^2^ of eQTL estimates in comparison to independent estimates from GTEx v7 [6].

## 3 Methods

### 3.1. Simulation Model

We assume we have access to reference panel data comprising *N* individuals with genotype matrix *X*, gene expression matrix *Z*, and case/control phenotype vector *Y*. The conventional method for calculating eQTLs involves a linear model over the observed *X* and *Z*. We also assume that we have a large GWAS data set comprising *N*′ >> *N* individuals with genotype matrix *X*′ and case/control labels for the same phenotype *Z*′. All SNPs and transcripts are normalized to mean 0 and standard deviation 1. We denote the set of SNP-gene linear effects *β*_1_ and the gene-trait linear effects *β*_2_. For our simulations, we oversimplify the genotypes to be sampled independently with an effect allele frequency set to 0.5 across all genes. We assume true gene expression quantities are distributed *Z* = *Xβ*_1_ + *ϵ* with 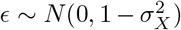, where 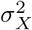 is the fraction of variance in *Z* explained by all predictors in *X*. The mediated SNP-trait effect is then their product *β*_1_*β*_2_, given all SNPs are sampled independently. For now, we assume there are no direct SNP-trait effects not mediated by gene expression. As *Y* is a binary variable, we opt to use a liability threshold model over the total SNP and gene effects 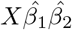, described in the next section.

The TWAS approach assumes first that an estimate 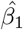 is calculated from the reference panel only, and then *β*_2_ is obtained by regressing *Y* against 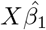. Therefore the accuracy of 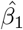 is one limitation of TWAS, as current sample sizes for gene expression data tend to be much smaller than for genotype data. Our goal is to perform the analysis in reverse: given the same data and an estimate of the TWAS effect size 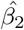, we would like to improve estimates of 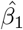. This relies on an assumption that 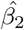, likely estimated initially from the same data, is robust to errors in 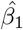 to a degree that allows its use to improve re-estimation of *β*_1_, provided that the gene in question mediates a significant fraction of the observed SNP-trait effects. Improving the accuracy of SNP-gene effects has the potential to improve TWAS estimates further through more accurate gene imputation. Additionally, re-estimating significance of eQTLs may provide more insights into regulation of gene expression by SNPs.

### 3.2. Liability Threshold Model

The liability threshold model is an alternative to logistic regression for identifying predictors (SNPs or genes) that are highly associated with disease status. This model assumes that the variance of all modeled predictors is a fraction of a normally distributed underlying quantitative liability, which is thresholded to generate case/control status. This model provides an advantage over the more conventional logistic model by explicitly modeling the variance explained, rather than estimating it by pseudo-*r*^2^ methods. It also provides a more natural interpretation of the Polygenic Risk Score (PRS).

Assume we are regressing measured or imputed quantitative gene expression values *Z* against case/control status *Y*. The liability threshold model assumes an underlying standard normal distribution representing the hidden liability. The PRS vector *Z*^*T*^ *β* explains a fraction of the variance of this liability, represented by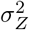. We describe in Section 3.3 a method to estimate 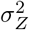 independently of the inference process for effect sizes in *β*. The liability is dichotomized into a sampled phenotype by labeling the sampled individual a case if the total liability score passes a threshold *T*, set such that 1 − Φ(*T*) is equal to the population prevalence if Φ(·) is the standard normal CDF. If an individual has polygenic risk score 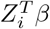, then the conditional mean of the liability is shifted by that value, and the variance of the liability is reduced by the variance of the PRS. Therefore, the probability of disease *Y*_*i*_ for an individual with predictors *Z*_*i*_ is given by Wray and Goddard [15]

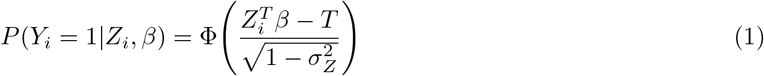

Our objective is to infer values of the coefficients *β* such that the probability of observed disease statuses across all individuals is maximized given their PRS. We assume the prior for all *β* to be a normal distribution with standard deviation 1. Therefore, the joint log likelihood of the liability threshold model is

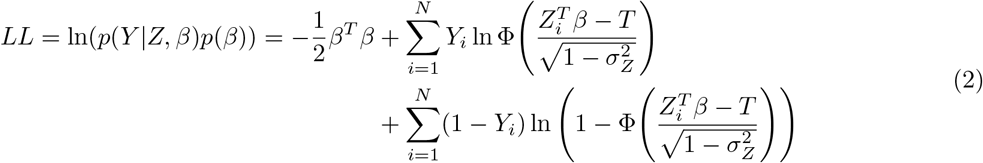

However, inference with respect to *β* is difficult because of the dependence of 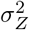 in the denominator on values of *β*.

### 3.3. Correction for effect size inflation

To resolve the dependence of 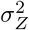 on *β* during inference, we instead independently estimate 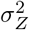 using the sample data prior to inference of *β* itself. A separate step to estimate total variance explained of predictors serves two roles: first, it decouples the PRS variance explained from the predictors themselves, facilitating inference, and second, it allows for correction of inflated effect size magnitudes inherent to both liability threshold model and logistic regression. Even when the relative magnitudes of effect sizes are accurately estimated, the overall scale of the effects is poorly controlled in both liability threshold and logistic models (see Results for simulations in which inflated effects produce larger likelihoods). By estimating the true total variance explained 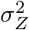, we can correct for this inflation by re-scaling all effect sizes by a constant factor such that the theoretical total variance explained as a function of *β* equals the observed 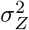.

When regressing predictors against a quantitative label, the total fraction of variance explained by the predictors can be obtained using Haseman-Elston regression [18, 19]. Golan et al. [20] demonstrate that a similar proportionality between labels and predictors applies to binary case/control labels in a liability threshold model. Assume the disease has a population prevalence *K*, study prevalence *P*, and liability threshold *T* = 1 − Φ(*K*) corresponding to the population prevalence. We can define a correlation between samples in *Y* by taking the outer product of all pairs (*i, j*) of standardized labels: a Bernoulli label *Y*_*i*_ is standardized as 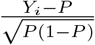. As in quantitative traits, this label correlation can be regressed against the correlation between pairs of samples in *Z*. Specifically, Golan et al. [20] show that correlations between labels and between predictors are related by the following linear function

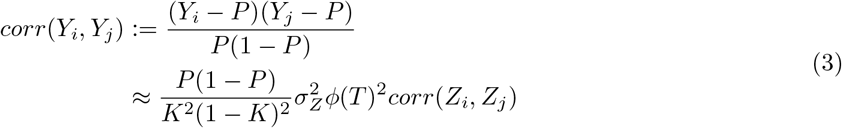

where 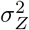 is the variance explained by *Z*. Therefore, we can regress the two sets of correlations over pairs of samples to obtain a slope, and then solve for 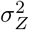 given observed population parameters of *P*, *K*, and *T*.

### 3.4. Expectation Maximization of the liability threshold model

By estimating a fixed value for 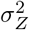, we have decoupled it from inference on *β*, resolving the aforementioned intractability. Therefore, we can perform inference by the conventional probit expectation maximization. First, we introduce a hidden variable *l* denoting the unobserved quantitative liability which determines case/control status *Y* = *P* (*l* > *T*). This results in the following EM equation for the joint log likelihood:

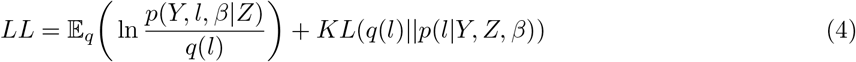

We assume *l* follows a standard normal distribution, with a fraction of variance explained by SNP effects: therefore 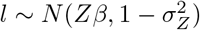. Furthermore, the distribution of *l*_*i*_ is dependent on disease label *Y*_*i*_, as *l* must fall on the correct side of the threshold *T* to impart the correct case/control status. Therefore, we have the following for the distribution *q*(*l*)

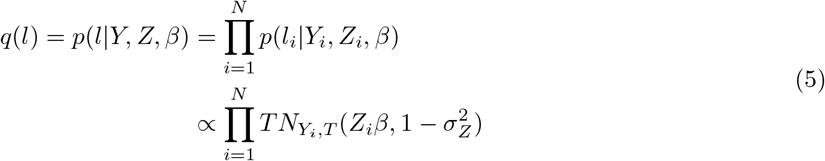

where *T N*_*Y*_ (·) is the truncated normal distribution over values greater than or less than *T* depending on whether *Y*_*i*_ indicates a case or control, respectively. We can then plug this distribution into the EM equation with joint log likelihood *LL*. By setting *q*(*l*) = *p*(*l|Y, Z, β*), the KL divergence term cancels, allowing us to maximize *β* with respect to the expected log likelihood over *q*(*l*). This term can be expanded as follows

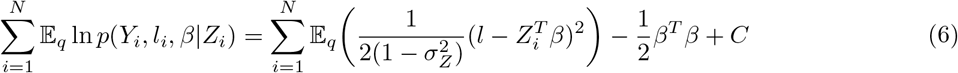

where *C* denotes terms which are constant with respect to *β*. Therefore, we have the following EM procedure, where in the E-step we update expected values of the hidden liability, and in the M-step we maximize *β* with respect to the likelihood informed by those expected liabilities.

**E-step**

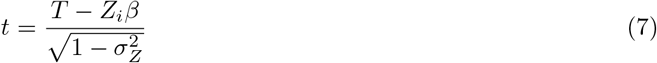

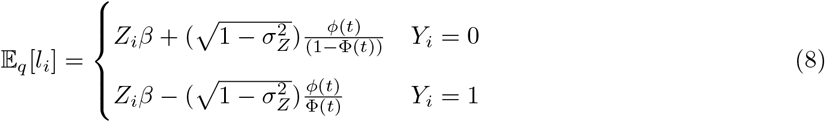
**M-step**

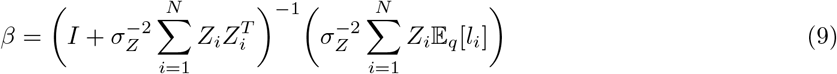 At each step the joint log likelihood is evaluated for convergence, after which the inferred *β* is returned.

#### 3.4.1. Constrained EM incorporating *h*^2^ correction

The method to estimate 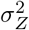 independently of *β* inference described by Golan et al. [20] allows us to correct for the aforementioned inflation inferred effect size magnitudes. As effect sizes tend to be scaled by an arbitrary constant factor, we can simply re-scale all effects such that their theoretical variance explained (*β*^*T*^ *β* if predictors in *Z* are standardized) equals 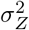 obtained by the Golan et al. [20] method.

However, we also investigated whether the re-scaling procedure can instead be incorporated directly into the EM algorithm as a constraint. Having obtained an estimate of 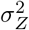 from the sample data, we add a Lagrangian multiplier to the maximization step to enforce the inequality constraint 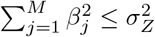. We now maximize the modified log likelihood

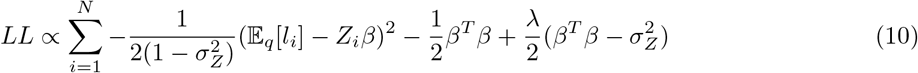

We notice that this modification conveniently adds only a constant term to the coefficient of *β*^*T*^ *β*.

Therefore, the new update equation has the following form

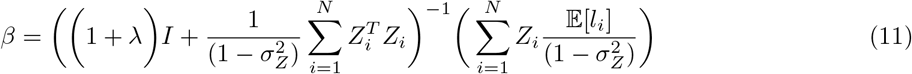

Note that the addition of lambda effectively re-scales the variance of the prior distribution of *β*, which has until now been assumed to be 1. At every iteration, we would like to solve for lambda such that the constraint 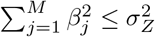 is satisfied. However, this operation does not appear to have an analytical solution, and we would like to limit the number of matrix inverse calculations required in a numeric solution. To accomplish this, we define the eigendecomposition for the covariance matrix 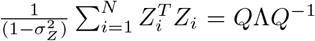. We also substitute *λ*′ = *λ* + 1. Therefore, the matrix inverse in equation 11 can be avoided by retaining the decomposition and only updating the diagonal matrix Λ, which requires only a vector division.

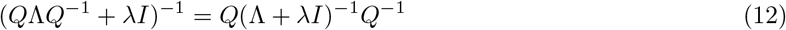

We can then express the predictors *β* in equation 11 in terms of their eigendecomposition, and plug in this term into the Lagrangian constraint.

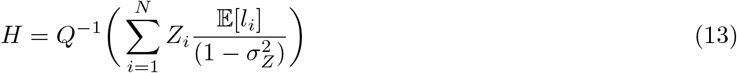

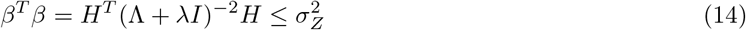

To our knowledge there is still no analytical solution for *λ* such that *β*^*T*^ *β* is close to 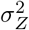. We instead take the numeric inverse to obtain 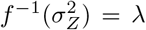. We perform this using the Python package pynverse, which performs a search over scalar values for *λ*. The operation in equation 13, though performed repeatedly, is relatively efficient as it can be performed as a dot product over 1-D vectors.

In the Supplementary Text, we have additionally derived an extension of the Lagrangian method for correlated predictors, in which case the variance of their weighted sum is dependent on the covariance between predictors.

### 3.5. Improving *β*_1_ estimation

Here we describe the main contribution of EMBER: a method to improve SNP-gene associations (*β*_1_) given an existing estimate of TWAS gene-trait effects (*β*_2_). Instead of a linear regression over the reference panel only, we combine this stage with a liability threshold regression passing estimates of *β*_1_ to scoring *X*′*β*_1_*β*_2_ against case/control status in a much larger GWAS cohort. We model one gene at a time, so *β*_2_ is a scalar value. First, we describe an adjust to the liability threshold model which accounts for increased error in imputed gene samples.

#### 3.5.1. Accounting for imputed gene expression in the liability model

To combine both observed expression *Z* and imputed expression *Z*′ = *X*′*β*_1_ into a single model, we introduce a hidden variable 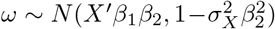 representing the distribution of the imputed gene effect on disease liability, where 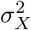 is the total fraction of variance in *Z* explained by predictors in *X*. We can now redefine disease risk with respect to *X*′ and *β*_1_ by marginalizing over the distribution of *ω*.

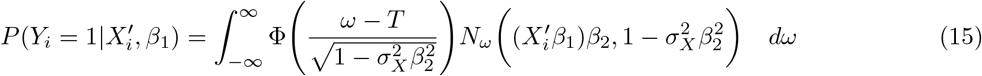

We can simplify this expression by applying the following lemma described by Owen [21]

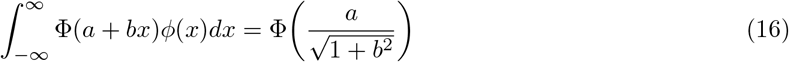

We define a standardized gene expression variable *ω*′ and substitute this into equation 15

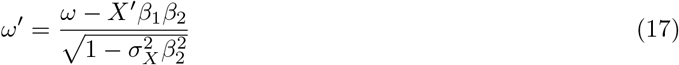

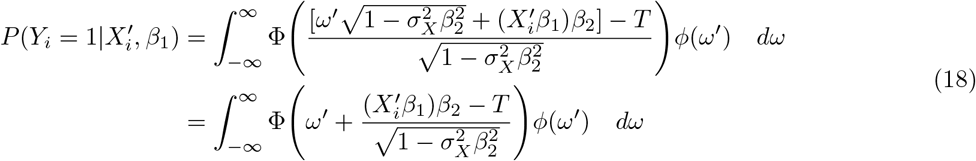

We can plug this expression into equation 16 with *b* = 1 and 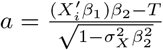, obtaining

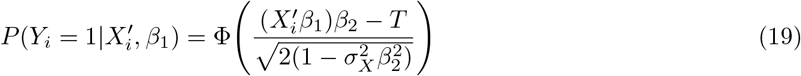

Therefore the only change from the liability model with observed *Z* instead of imputed *X*′*β*_1_ is the non-explained liability variance is multiplied by a factor of 2. This distinction in variance explained allows us to perform regression on both observed gene expression in the reference panel and imputed gene expression from GWAS simultaneously.

#### 3.5.2. Inference incorporating both observed and imputed gene expression

Our goal is to infer SNP-gene effects given access to both observed genotype and gene expression data from a small reference panel as well as genotype and trait data only from a large GWAS cohort. Therefore, the log joint likelihood now comprises a sum between the normal prior on *β*, a linear regression over observed gene expression in the reference panel, and liability threshold regression over imputed gene expression in a GWAS cohort. We first estimate 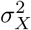, the variance explained of SNPs on gene expression, using either H-E regression on the reference panel or the liability method by Golan et al. [20], making sure to divide by 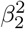 to remove the contribution of the gene-trait effect. The joint log likelihood can then be expressed as

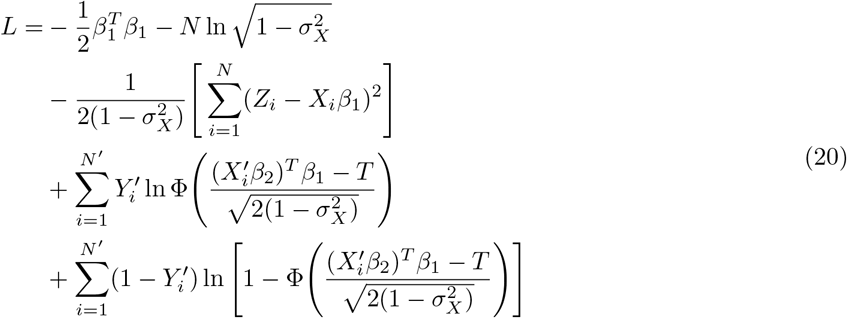

The inclusion of linear regression terms can be accommodated easily in the EM algorithm. The new updates are

**E step:** Calculate 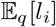 with 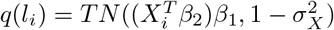

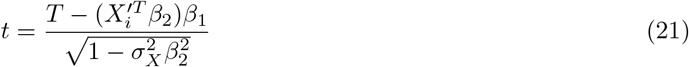

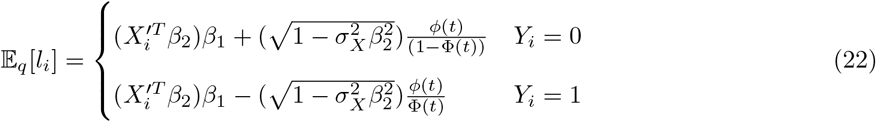
**M step:**

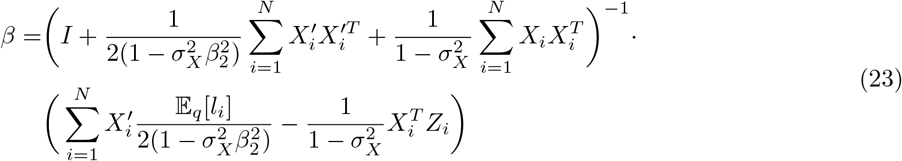

The inferred *β* can be corrected for effect size inflation by either re-scaling according to 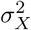 after EM inference, or since the form of the EM algorithm is largely unchanged, the Lagrangian can be incorporated as described in section 3.4.1.

## 4 Results

### 4.1. Robustness of TWAS estimates (*β*_2_) to sample size

Broadly, we are interested in whether simultaneous inference of SNP-gene (*β*_1_) and gene-trait (*β*_2_) effect sizes can improve accuracy over two-stage regression, given access to both a reference panel and an independent GWAS cohort. As GWAS cohorts are often orders of magnitude larger, hidden gene expression quantities must be inferred for a large majority of the data. The accuracy of these hidden values, which rely on estimates of *β*_1_, may in turn improve estimation of *β*_2_. But to improve *β*_1_ estimation over an initial result from the observed reference panel data, we must incorporate the GWAS cohort and our best estimate of *β*_2_. As the potential benefit of EMBER derives from providing additional samples with only partially observed data, we first investigate the robustness of TWAS associations to sample size. While Gusev et al. [11] report that increasing the size of reference panels past several hundred samples does not improve eQTL estimation, Wainberg et al. [14] nevertheless claim more recently that dependence on small eQTL reference panels is a potential limitation for TWAS.

We found in simulations that TWAS-imputed gene-trait effects are very robust to small sample size in the reference panel, corroborating Gusev et al. [11], and therefore we do not expect significant improvement in *β*_2_ estimation from combining reference and GWAS data sets. We ran simulations comparing *r*^2^ of inferred results from a probit regression to the true generating effects. These results are shown in Figure S1 as a function of sample size and varying expression-trait variance explained. We find across all conditions that the performance of TWAS plateaus at an *r*^2^ of 0.8, and this value is reached at a reference panel size of roughly 500, similar to that of the CommonMind Consortium (CMC) data, for all values of gene-mediated variance explained except for the smallest of 0.01. These results indicate that TWAS is quite robust to sampling error due to small reference panel sizes, and there is little room for improvement on this front. This also suggests that an estimate of *β*_2_ from data is sufficient to improve estimation of eQTLs (*β*_1_) in one step, rather than requiring repeated updates of one set of effects given newer estimates of the other.

### 4.2. Constrained probit EM for correct effect sizes

We demonstrate that effect size inflation does occur in conventional applications of logistic and probit regression by simulating case/control data with a single predictor and known effect size. From generated case/control samples, we then calculate the log likelihood resulting from scaling the known effect size by a scalar factor (Fig S2A). We find that the scalar factor resulting in the maximum log likelihood does not occur at a value of 1.0, corresponding to the original value, but rather at roughly 1.4, indicating a significant inflation from the inference.

By simulating case/control data from a probit/liability threshold model, we observe inflation in the inferred effects of both the logistic and probit models in Fig S2B. While the log odds ratios of the logistic model are not expected to map directly onto liability model used to simulate the data, we should expect to recover effects of the correct magnitude in the probit model. We rescale the effects by calculating the total variance explained of the predictors as described in the Methods and then divide each effect size by a constant such that their sum of squares equals the calculated value. Lastly, the faster constrained EM method produces nearly identical estimates to those of the slower rescaling method (Fig S2B).

To demonstrate the performance advantages of the constrained EM algorithm in comparison to rescaling the conventional liability threshold model, we measured performance time and number of EM iterations as a function of increasing sample size with other parameters kept constant. Results are shown in Fig 2, where for both methods the number of iterations remains relatively constant as a function of sample size, and the total execution time increases linearly. By both metrics, the constrained probit method far outperforms the rescaling method by requiring fewer iterations despite performing more work per iteration.

**Figure 1:**
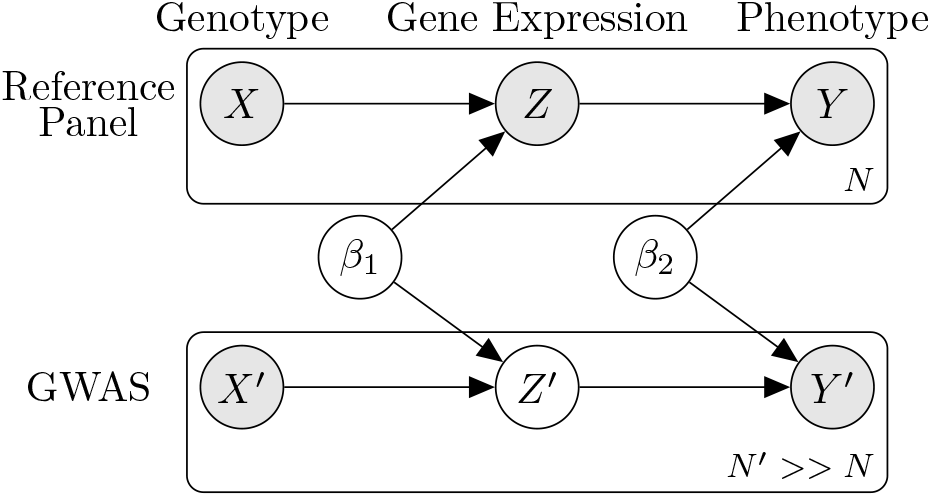
For a small reference panel cohort of size *N*, we observe SNPs *X*, transcripts *Z*, and binary trait *Y*. Assume we have a large GWAS cohort of size *N*′ with only *X*′ and *Y*′ observed. EMBER attempts to improve inference of *β*_1_ (the SNP-gene effects) given *β*_2_ (the TWAS gene-trait effect).

**Figure 2:**
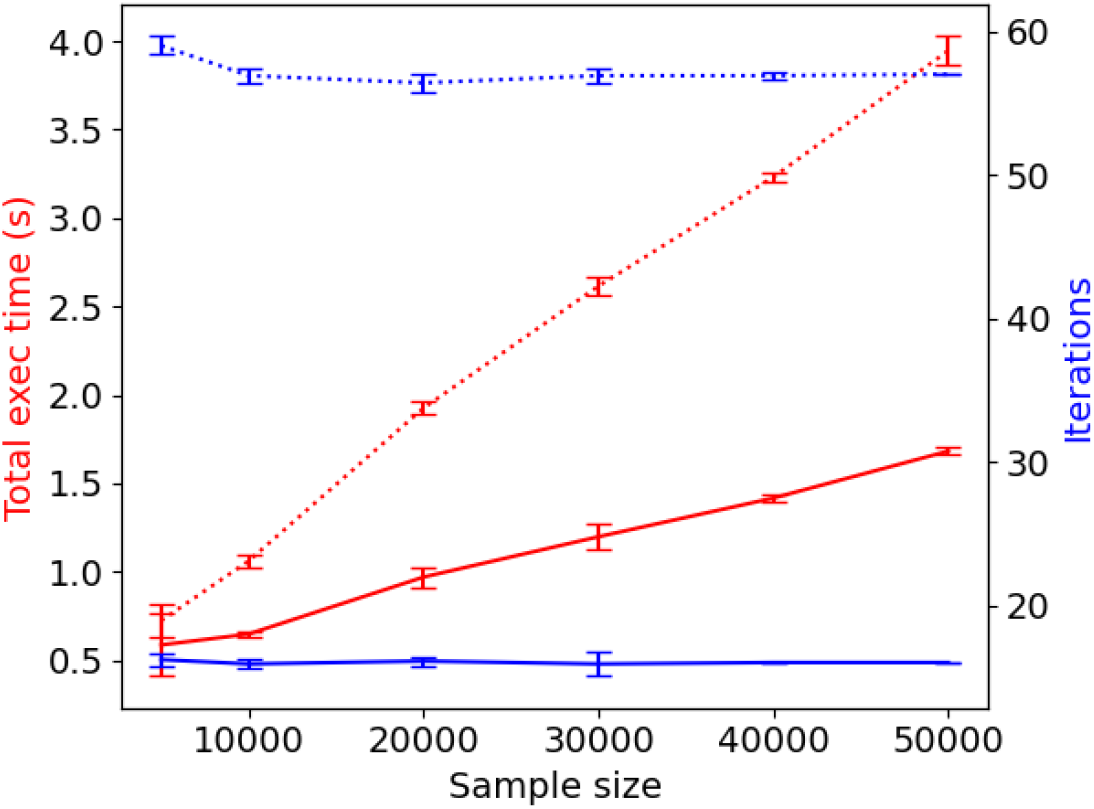
Performance of constrained probit model (solid line) versus conventional probit with rescaling (dotted line). Performance was measured as a function of sample size in terms of total execution time (red) and total number of iterations to convergence (blue). All simulations are run with 200 standard normal predictors with a total variance explained of 0.1. The mean and standard deviation of 10 trials is shown for each sample size.

### 4.3. Reversing TWAS to re-estimate eQTLs

To test whether we can improve estimation of SNP-expression effects (*β*_1_) when a reference panel and GWAS cohorts are analyzed together, we ran the EMBER method for estimating eQTLs described in the Methods on simulated data. We simulated sample sizes roughly corresponding to those of the CMC and PGC data: a reference panel size of 500 and GWAS size of 50,000. We also assumed for now that the variance explained of non-gene mediated SNP effects is zero. An example run is shown in Fig 3. In this single trial, the *r*^2^ of EMBER more than doubles that of linear regression: 0.88863 versus 0.30665. Additionally, EMBER correctly re-scales all effects so that their magnitudes fall appropriately on the *y* = *x* line.

**Figure 3:**
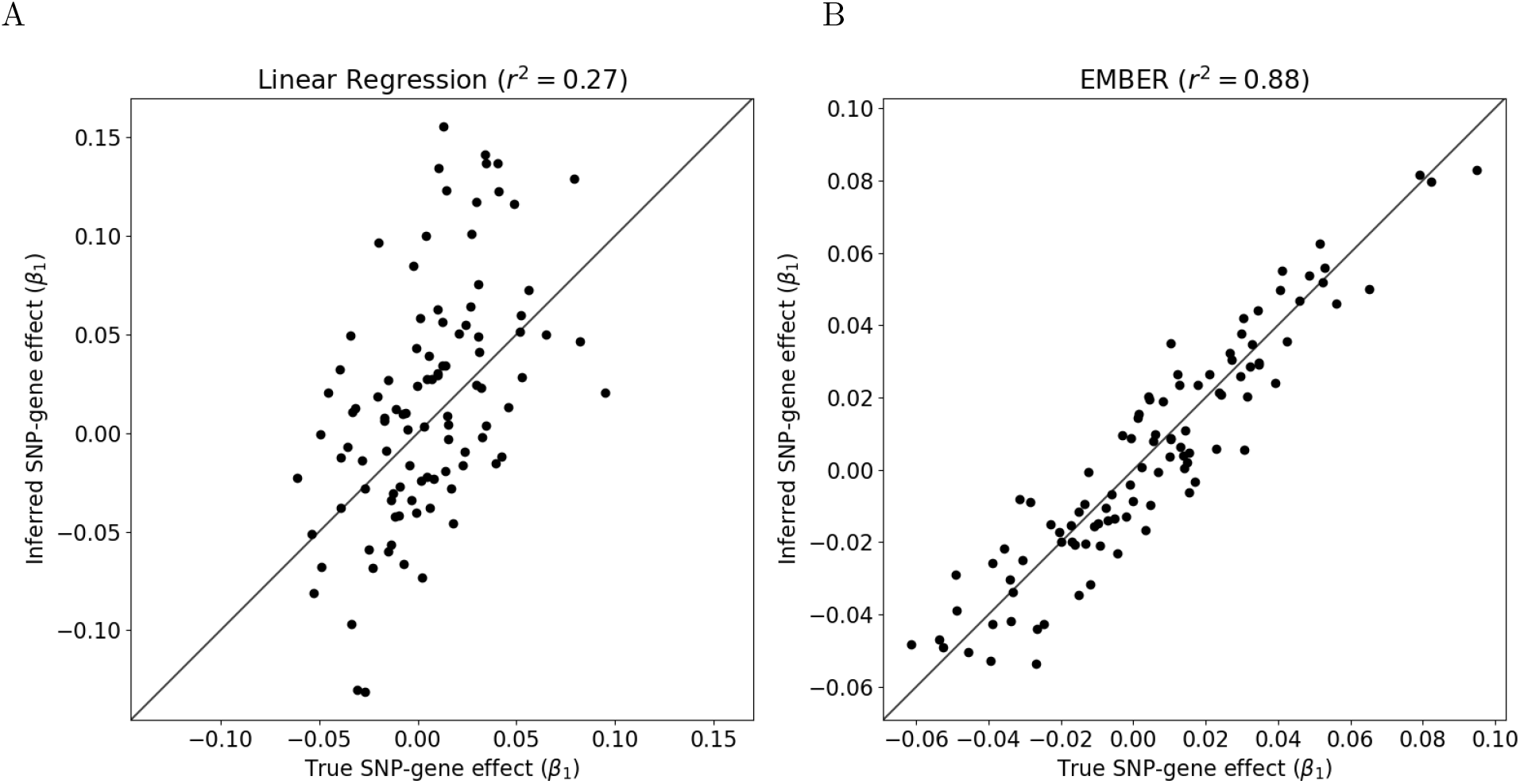
Inference of eQTLs on simulated data by (A) conventional linear regression on the reference panel only and (B) combining reference panel and GWAS data in EMBER. For simulation we assume a reference panel size of 500, GWAS sample size of 50,000, 100 SNPs and one gene with total SNP-gene variance explained of 0.1 and gene-trait variance explained of 0.01. A population prevalence of 0.01 was assumed and allele frequency of 0.5 for all SNPs.

EMBER assumes that there are no non-gene mediated SNP effects, but in practice, this assumption may be violated. To test this, we fixed the gene-mediated SNP variance explained to a constant, and varied the non-mediated variance as a percentage of that constant. These results are shown in Fig 4. We observe that when the percentage on the x-axis is zero, corresponding to results shown in Fig 3, EMBER significantly outperforms linear regression to a mediated variance explained of 0.005, after which the expression layer contributes very little signal to the inference. We also observe that even though the assumption of no non-mediated effects is violated, EMBER still outperforms linear regression up to a percentage of non-mediated variance between 0.7 and 0.8 across the tested conditions.

**Figure 4:**
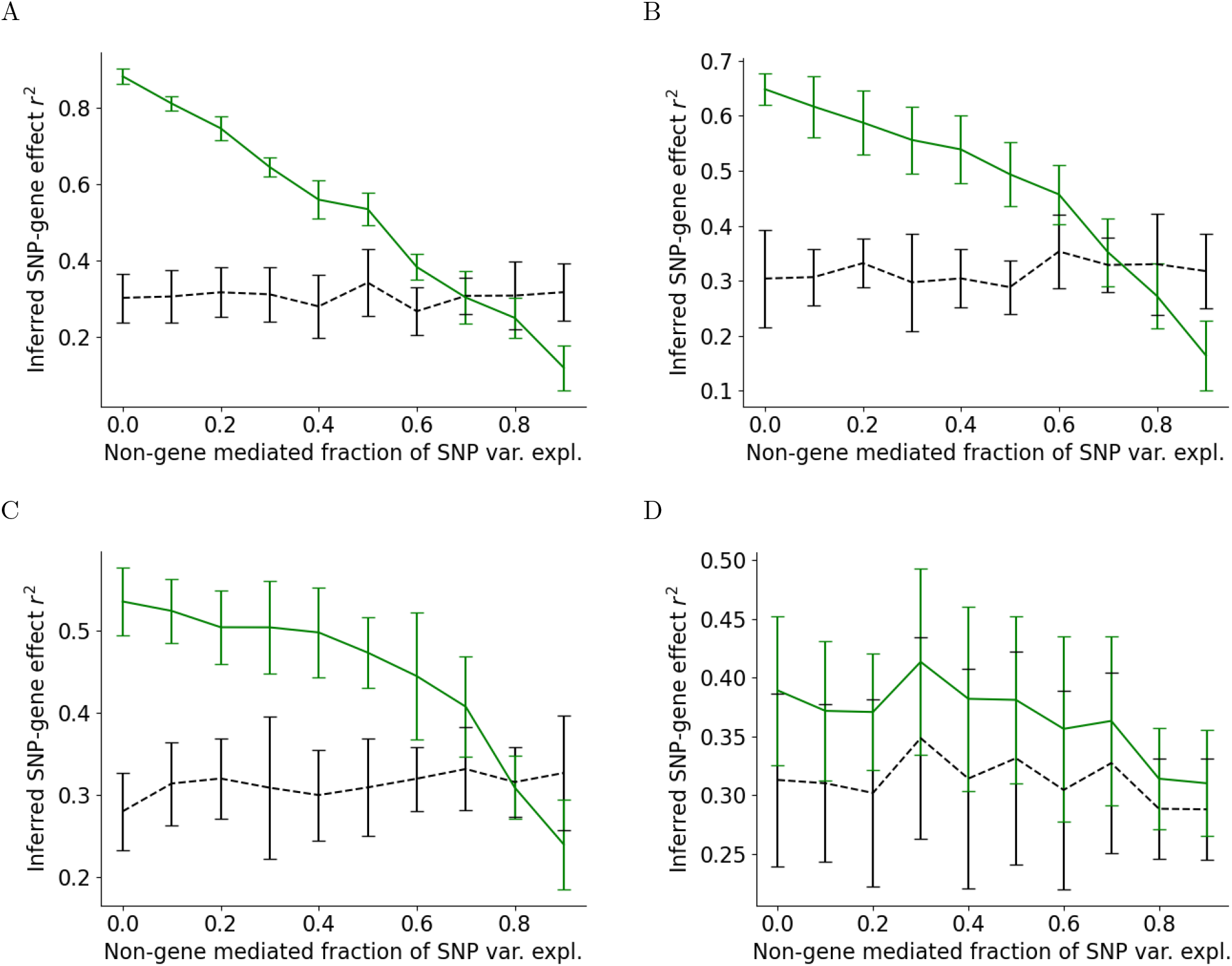
EQTL estimation (*r*^2^) as an increasing fraction of non-mediated SNP variance explained for EMBER (green) and linear regression (black). The gene-mediated SNP variance explained is fixed to (A) 0.05, (B) 0.01, (C) 0.005, (D) 0.001. Percentages are specified as 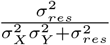 where 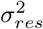 denotes the non-gene mediated residual SNP variance explained, and 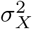 and 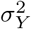 denote the SNP-gene and gene-trait variances explained by *β*_1_ and *β*_2_, respectively.

### 4.4. Validation in schizophrenia

We applied EMBER to study eQTLs for genes relevant to schizophrenia. We obtained a gene expression reference panel from the CommonMind Consortium (CMC) and GWAS data from 34 cohorts of the Psychiatric Genomics Consortium (PGC). For quality control, we followed the suggestions in Marees et al. [22]: we removed individuals with genotype missingness rate greater than 0.05, and we removed SNPs with missingness rate in the data greater than 0.1, minor allele frequency less than 0.01, and Hardy-Weinberg equilibrium less than 1 × 10^−10^. We additionally filtered for individuals in the CMC data with both genotype data and expression data available. This preprocessing resulted in 512 individuals from the CMC data and 56,081 individuals from the PGC. We focused on genes which have been shown to be associated with schizophrenia in TWAS by Mancuso et al. [13], obtaining 36 genes with TWAS p-value < 10^−5^ and which are also present in the CommonMind data. We imputed cis-SNPs using SHAPEIT [23] and Impute2 [24, 25] and obtained ancestry principal components from PLINK [26]. As EMBER is dependent on gene mediation to improve performance, we calculated the fraction of SNP variance mediated by expression by separately calculating total SNP variance, and SNP-expression variance multiplied by inferred TWAS effect squared. These results are shown in Fig S3A. To obtain the best possible TWAS effect sizes for our data, we re-calculated them in the usual manner by estimating eQTLs from the CMC reference data only, imputing gene expression for the PGC data, and then regressing imputed expression against case/control status. Covariates included for all analyses were sex and top 10 ancestry PCs.

To validate EMBER, we performed eQTL re-estimation in both EMBER and linear regression on the CMC and PGC data, and compared the level of concordance with reported eQTLs in GTEx v7 data for the frontal lobe. A high degree of concordance would suggest reproducibility of effects and a higher confidence that these are true eQTLs. We obtained GTEx v7 eQTLs effects from the TWAS/Fusion website (http://gusevlab.org/projects/fusion/), calculated using BLUP.

To sidestep challenges incorporating the model assumptions of EMBER into the BLUP method, we instead addressed significant LD in cis-SNP regions by performing dimensionality reduction over the input SNPs and covariates by performing a second PCA over all predictors, not to be confused with the initial PCA used to discover ancestry covariates. After inferring effects on the transformed PC variables, the predictor-level effects can be recovered by multiplying the PC loadings matrix by the inferred effects over PCs. As the number of components to include from the variable reduction PCA is an open question, we decided to test the performance of linear regression results compared to GTEx reported effects as a function of this parameter. The mean *r*^2^ compared to GTEx across all 36 genes is shown in Fig S3B as a function of number of PCs of the variable reduction PCA to ensure independence of input predictors. We observe that the performance plateaus between 5 and 10 PCs and begins to decline afterward, which prompted us to select 10 PCs for this procedure.

We evaluated the accuracy of baseline linear regression and EMBER, both using the variable reduction preprocessing step, by the concordance of their inferred SNP-expression effects with those reported by TWAS/FUSION in GTEx v7 data. As TWAS/FUSION also reports effect size calculations from the CMC data for 26 out of the 36 studied genes, we compare the *r*^2^ of these results against the GTEx effects when possible, along with our own implementation of the BLUP method for all genes. We assume in both the EMBER and BLUP models a normal prior effect size distribution with a variance of 1 × 10^−4^. Results for these methods across all 36 genes are shown in Figure 5. Across all genes, EMBER achieves an average *r*^2^ of 0.235 ± 0.145, improving over an *r*^2^ of 0.200 ± 0.129 obtained by linear regression. EMBER also improves *r*^2^ in 26 out of 36 genes (Figure 5B), with a one-sided sign test p-value of .00197. However, we do not observe a significant relationship between improvement in performance and the fraction of gene-mediated variance explained in Figure 5B. Given the generally low fraction of variance explained across all genes (see Figure 4), it is possible that cis-SNP effects not mediated by the studied genes contribute to a reduction in estimation accuracy. The average *r*^2^ achieved by EMBER is slightly lower that of our implementation of the BLUP model at 0.240 ± 0.123 (Figure 5A), though both methods perform significantly better than the same BLUP model on CMC data as reported by TWAS/FUSION, with an *r*^2^ of 0.201 ± 0.109. As EMBER is currently applied to a baseline linear regression model with a variable reduction step, further improvements in performance may be possible by accessing the full SNP covariance matrix as in the BLUP method. Figure 6 displays effect size comparisons for the gene GLT8D1, where we observed the largest increase in *r*^2^ in EMBER compared to linear regression. Additional plots for all genes are shown in Figure S4.

**Figure 5:**
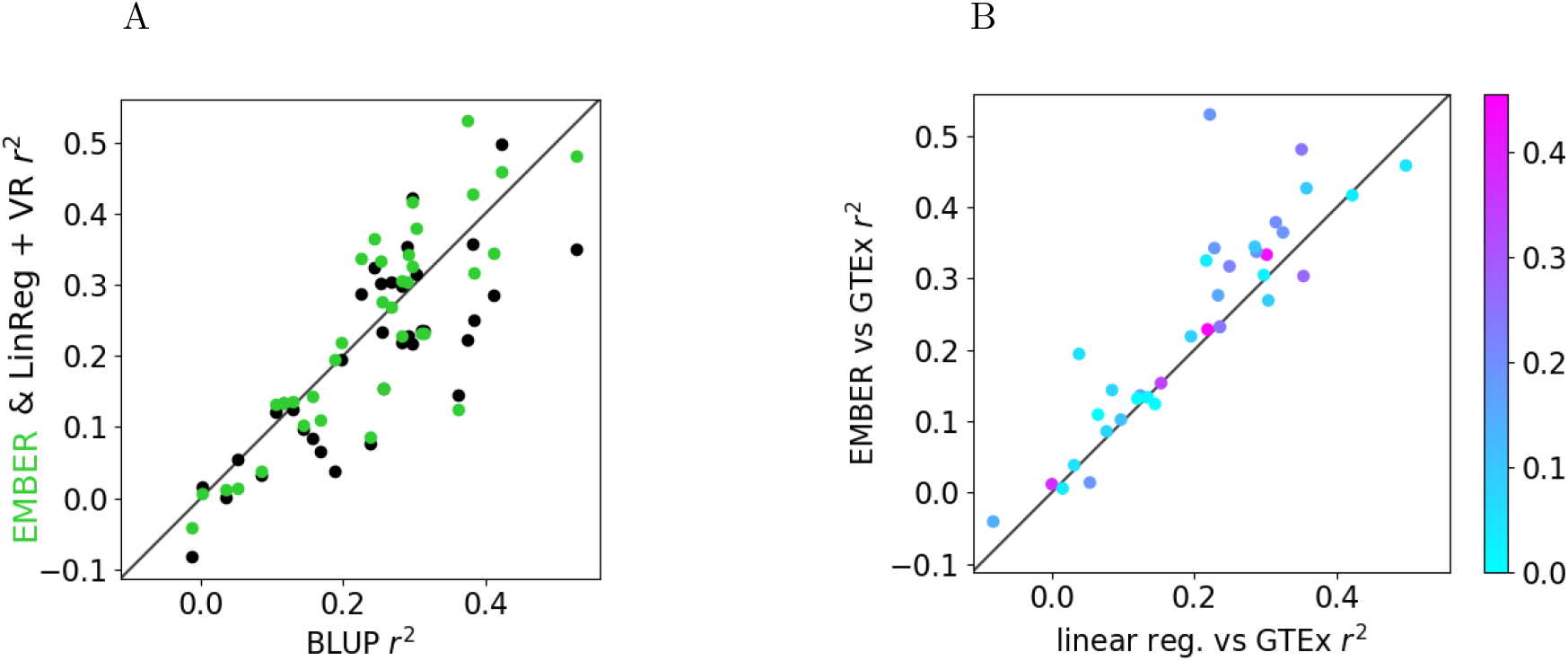
(A) For 36 genes, concordance of estimated SNP-expression effects with reported GTEx v7 effects using BLUP versus using EMBER (green) and linear regression (black) with a variable reduction step. (B) Concordance with reported GTEx v7 effects of linear regression versus EMBER. Colors indicate fraction of SNP variance explained that is mediated by gene expression.

**Figure 6:**
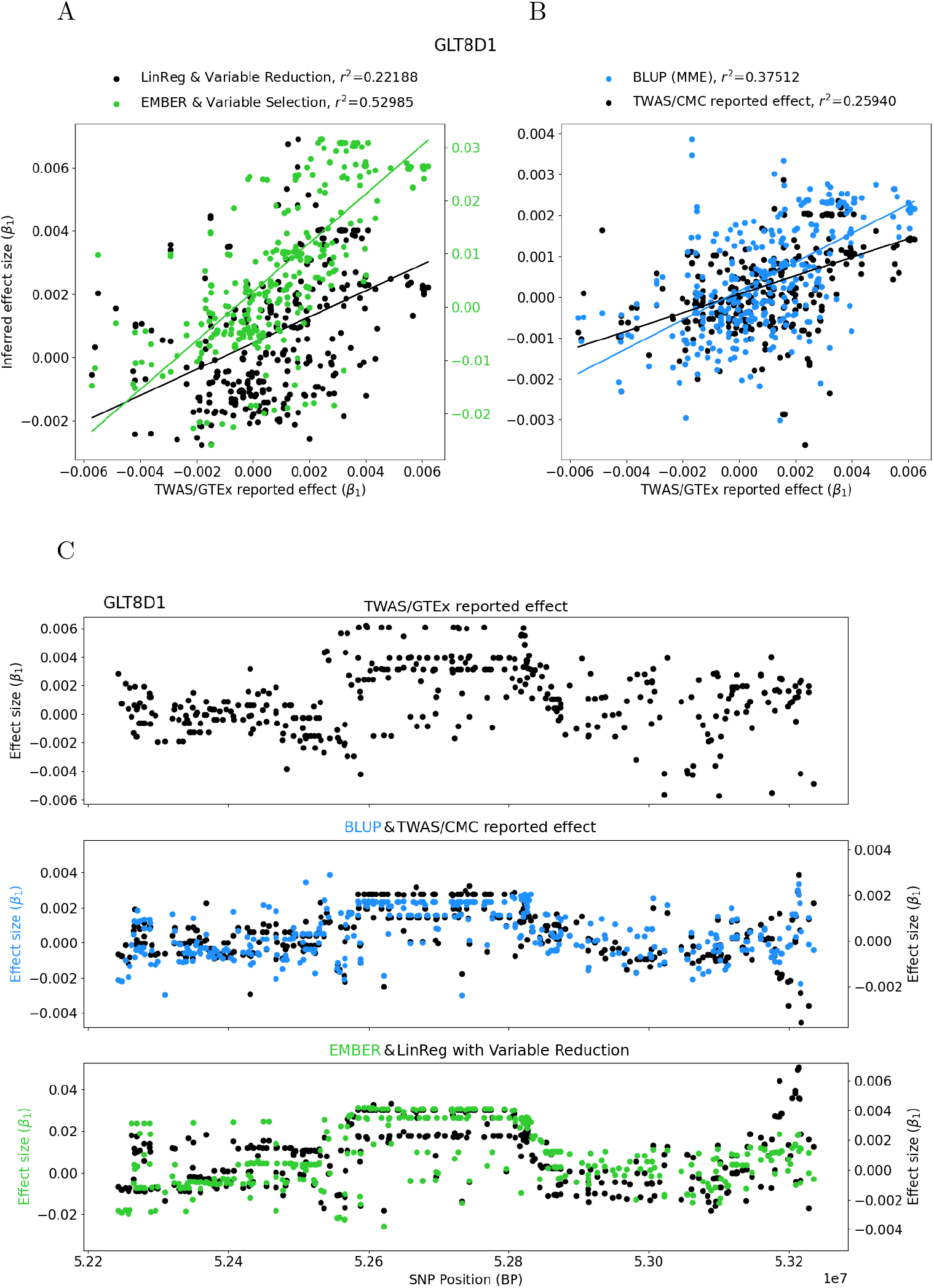
Example results for the gene GLT8D1, for which we observe the largest improvement in *r*^2^ using EMBER. (A) GTEx v7 effects reported by TWAS/FUSION versus effects inferred from linear regression (black) and EMBER (green) with variable reduction. (B) GTEx v7 effects reported by TWAS/FUSION versus effects inferred from BLUP (blue), as well as BLUP effects reported by TWAS/FUSION on CMC data (black). (C) Effect sizes for all models arranged by SNP position in the locus. The top row displays TWAS/FUSION reported BLUP effects on GTEx v7 data; the middle row displays BLUP effects (blue) and TWAS/FUSION reported BLUP effects on CMC data (black); and the bottom row displays results of EMBER (green) and linear regression (black), both using variable reduction to resolve high LD across SNP predictors. For all tested genes, see Fig S4.

## 5 Discussion

We presented EMBER, a method to perform inference over gene expression reference panel data alongside GWAS data simultaneously to improve estimation of effect sizes over a two-stage regression approach. In developing EMBER, we introduced also a variance correction to incorporate both measured and imputed gene expression as predictors into a single regression, and a constrained EM method to rapidly re-scale effect sizes based on expected variance explained.

We mapped the parameter space where EMBER is effective. Through simulations, we found that imputed gene-trait associations are quite robust to low reference panel sample sizes, but that significant improvements in eQTL estimates are possible by including GWAS data with no gene expression data observed. Additionally, while EMBER focuses on gene mediated effects, we found that accuracy of inferred effects still improved when the fraction of gene-mediated variance explained was as low as 0.3.

We observed significant improvement in concordance of inferred eQTL effects when we performed a variable reduction step involving transformation of predictors to lower dimensions by PCA. This allows us to account for significant multicollinearity resulting from LD. With this step, ordinary least squares regression achieves an *r*^2^ performance nearly equivalent to BLUP effects reported by TWAS/FUSION on CMC data. This allowed us to build EMBER from a linear regression model, although further improvements may be possible if more sophisticated multi-SNP models such as BLUP can be incorporated. These designs may present additional challenges: for example, a model combining EMBER and BLUP would require features from both the probit and linear mixed models, and while such algorithm have been proposed [27, 28], these typically require performing operations on the full covariance matrix of random effects, which is increasingly impractical for large GWAS.

EMBER is not without its limitations. After variable reduction, for several genes the improvement observed in CMC and PGC data is quite small. This is likely because the fraction of variance explained for these genes is lower than the threshold of 0.2-0.3 for gene-mediated fraction of SNP variance, and EMBER is not benchmarked to improve results in this range. Even so, we do not observe significant reductions in performance in EMBER relative to linear regression, with the largest drop in *r*^2^ being −0.05 in the gene MAPK3.

Going forward, EMBER demonstrates a more general approach to the investigation of genotype-to-phenotype association. Instead of relying on a single study with its traded-off weaknesses of sample size vs. extensiveness of phenotyping, we show that one can leverage modern inference tools in order to get the best of both worlds across multiple studies of different designs. In the era of ubiquitous availability of genotype data, this can greatly facilitate the discovery of new signals.

## 6 Web Resources

A Github page for EMBER, including code to reproduce figures is available at: https://github.com/jyuan1322/EMBER

## Notes

https://github.com/jyuan1322/EMBER

### Competing Interest Statement

The authors have declared no competing interest.

