## Supplemental Text and Figures for "Reversing Transcriptome-Wide Association Studies to improve expression Quantitative Trait Loci associations"

#### 7 Scaling of effects with correlated predictors

In our results a variable reduction step using PCA guarantees independence of input predictors. But in other applications, without this step correlatedness between predictors in a linear model will alter their total variance explained, so we must account for these correlations in order to produce the correct re-scaling of effect sizes. Like other regression models, we however cannot completely resolve multicollinearity from this correlatedness and must rely on large sample sizes to compensate.

When predictors are independent and normally distributed, the total variance explained of the predictors is the sum of squares of the effect sizes.

$$\sum_{j=1}^{M} \beta_j^2 = \beta^T \beta = \sigma_Y^2$$

Assume instead that predictors are sampled from a multivariate normal distribution with mean 0 and covariance $V$. In practice, we need to obtain $V$ from the sample covariance of controls. The total variance explained of the weighted sum of predictors is

$$\sum_{j=1}^{M} \sum_{k=1}^{M} \beta_j \beta_k \; cov(Y_j, Y_k) = \beta^T V \beta = \sigma_Y^2$$

If we have estimated $V$ from a control cohort, we can obtain a scaling factor $a$ such that $\frac{1}{a^2}\beta^T V \beta = \sigma_Y^2$, which rescales the effect sizes to the correct magnitude $\frac{\beta}{a}$ as before.

This however requires a change to the fast Lagrangian method. The new likelihood is

$$LL \propto \sum_{i=1}^{N} -\frac{1}{2(1-\sigma_Y^2)} (\mathbb{E}_q[l_i] - Y_i \beta)^2 - \frac{1}{2}\beta^T \beta + \frac{\lambda}{2}(\beta^T V \beta - \sigma_Y^2) \tag{24}$$

$$\beta = \left( I + \lambda V + \frac{1}{(1-\sigma_Y^2)} \sum_{i=1}^{N} Y_i^T Y_i \right)^{-1} \left( \sum_{i=1}^{N} Y_i \frac{\mathbb{E}[l_i]}{(1-\sigma_Y^2)} \right) \tag{25}$$

Because $V$ is not a diagonal matrix, we cannot use the eigendecomposition trick as before. However, we can apply the following theorem (using Neumann series representation for inverse):

$$\begin{aligned}(A+\lambda V)^{-1} = A^{-1} &- \lambda A^{-1}VA^{-1} \\ &+ \lambda^2 A^{-1}VA^{-1}VA^{-1} \\ &- ...\end{aligned} \tag{26}$$

We reduce computation time of $\beta^T V\beta$ by calculating only half of the symmetric product and then taking the sum of squares of the resulting vector. If we define the Cholesky decomposition for $V$,

$$\begin{aligned}V &= LL^T \\ C &= \sum_{i=1}^{N} Y_i \frac{\mathbb{E}[l_i]}{(1-\sigma_Y^2)} \\ A &= I + \frac{1}{(1-\sigma_Y^2)} \sum_{i=1}^{N} Y_i^T Y_i \\ \beta^T V\beta &= SumSquares\Big(C(A^{-1}L - \lambda A^{-1}VA^{-1}L)\Big)\end{aligned}$$

We now must take the sum of squares where each term being summed has the form $(r - r'\lambda)$. Taken across all terms we have the following

$$\begin{aligned}(r-r'\lambda)^2 + (s-s'\lambda)^2 + (t-t'\lambda)^2 = &r^2 \quad - \quad 2rr'\lambda \quad + \quad r'^2\lambda^2 \\ &s^2 \quad - \quad 2ss'\lambda \quad + \quad s'^2\lambda^2 \\ &t^2 \quad - \quad 2tt'\lambda \quad + \quad t'^2\lambda^2\end{aligned} \tag{27}$$

The solution for $\lambda$ is then the result of the quadratic equation with $a = r^2 + s^2 + t^2 + ...$, $b = -2(rr' + ss' + tt')$, and $c = r'^2 + s'^2 + t'^2 + ....$ In the above, $r = CA^{-1}$ and $r' = CA^{-1}VA^{-1}L$.

455 # 8 Additional figures and validation in CMC/PGC cohorts

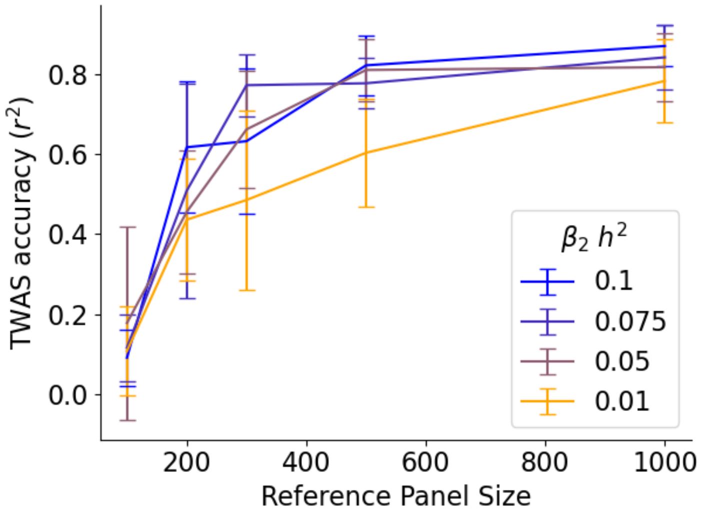


Figure S1: Accuracy of TWAS inferred gene-trait effects ($\beta_2$) as a function of reference panel sample size for 20 simulated genes. A set of SNP-gene effects is learned from this reference panel, and inferred genes are calculated for a fixed GWAS sample size of 50000 and regressed against the phenotype using a probit model. Colors indicate the total variance explained by the genes. The mean and standard deviation of 20 trials is shown. All simulations used a SNP-expression variance explained of 0.1.

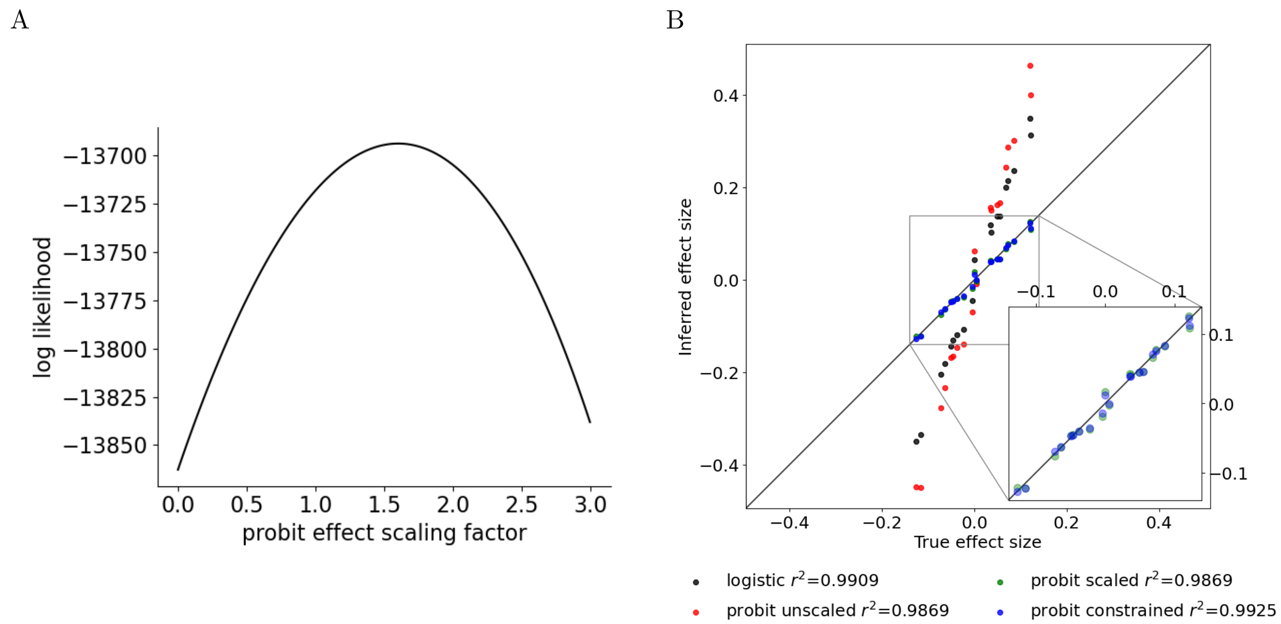


Figure S2: A. Effect size inflation in the true liability threshold model. A liability threshold model with a single predictor and known effect size was used to simulate case/control data. The log likelihood was then tested using the same effect size scaled by a constant on the x-axis, with 1.0 equivalent to the true effect. The maximum log likelihood occurs at a value larger than 1.0, causing inflation in the inferred effect size. B. True versus inferred effect sizes for simulations using 20 standard-normally distributed predictors (genes) with a total variance explained of 0.1, sample size of 5,000, and case/control prevalence of 0.01. While all models correctly infer the relative magnitudes of effect sizes, the logistic and liability threshold (probit) models by default over-estimate the scale of effects. By rescaling the effect sizes according to observed variance explained, we are able to correct for this inflation. The constrained method, which is more time efficient, produces nearly identical results to the post-inference rescaling.

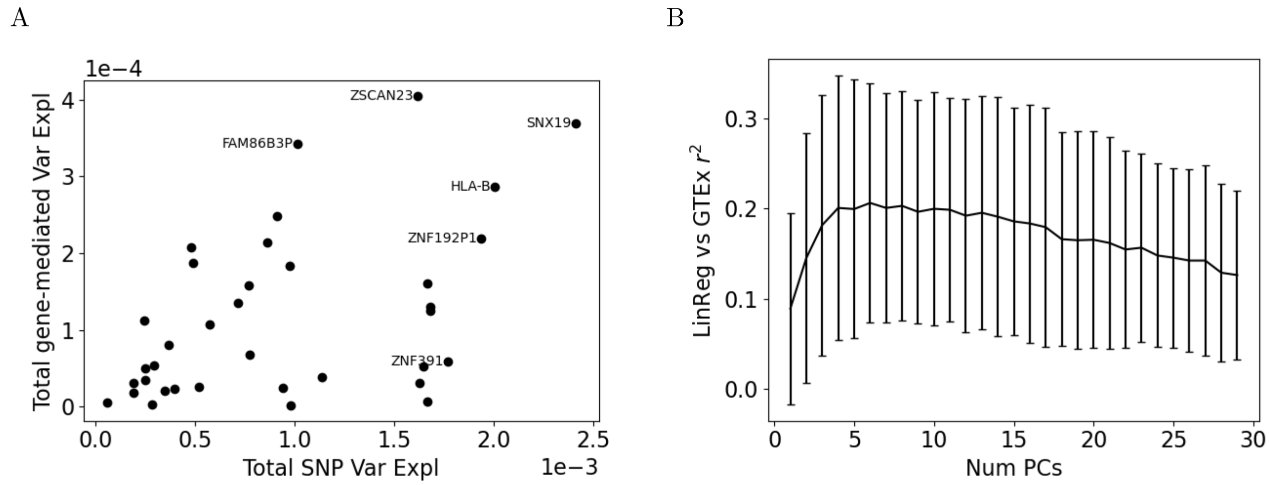


Figure S3: **A.** For the 36 genes with TWAS p-value $< 10^{-5}$, plotted are the total SNP variance explained against the fraction of SNP variance that is mediated by gene expression. **B.** Linear regression $r^2$ compared to GTEx v7 effects as a function of number of PCs in PCA for variable reduction over cis-SNPs.

| Gene | EMBER/GTEx $r^2$ | LR/GTEx $r^2$ | $\Delta r^2$ | frac mediated $h^2$ |
|---|---|---|---|---|
| GLT8D1 | 0.530 | 0.222 | 0.308 | 0.189 |
| DUS2 | 0.194 | 0.039 | 0.156 | 0.060 |
| ZSCAN23 | 0.481 | 0.351 | 0.130 | 0.250 |
| C5orf63 | 0.343 | 0.228 | 0.114 | 0.180 |
| ZNF391 | 0.325 | 0.217 | 0.108 | 0.033 |
| SMIM8 | 0.427 | 0.357 | 0.069 | 0.093 |
| PRMT7 | 0.317 | 0.249 | 0.068 | 0.219 |
| NAGA | 0.379 | 0.314 | 0.064 | 0.205 |
| BTN3A2 | 0.345 | 0.285 | 0.060 | 0.097 |
| ZKSCAN3 | 0.144 | 0.085 | 0.059 | 0.075 |
| RNASEH2C | 0.338 | 0.287 | 0.050 | 0.195 |
| BTN2A1 | 0.109 | 0.065 | 0.044 | 0.004 |
| SNHG5 | 0.277 | 0.233 | 0.043 | 0.160 |
| HLA-B | -0.041 | -0.082 | 0.041 | 0.143 |
| CORO7 | 0.365 | 0.324 | 0.040 | 0.187 |
| GIGYF1 | 0.333 | 0.302 | 0.032 | 0.431 |
| ARL14EP | 0.219 | 0.195 | 0.023 | 0.087 |
| SNX19 | 0.136 | 0.124 | 0.012 | 0.152 |
| GNL3 | 0.132 | 0.120 | 0.012 | 0.002 |
| TOM1L2 | 0.012 | 0.000 | 0.011 | 0.379 |
| ZNF738 | 0.229 | 0.219 | 0.010 | 0.455 |
| ZSCAN31 | 0.086 | 0.077 | 0.009 | 0.077 |
| PPM1M | 0.305 | 0.298 | 0.007 | 0.025 |
| FOXN2 | 0.039 | 0.032 | 0.007 | 0.058 |
| ZNF192P1 | 0.102 | 0.097 | 0.005 | 0.113 |
| FAM86B3P | 0.153 | 0.153 | 0.000 | 0.338 |
| XPNPEP3 | 0.134 | 0.135 | -0.001 | 0.019 |
| AS3MT | 0.233 | 0.236 | -0.003 | 0.247 |
| CBR3 | 0.232 | 0.236 | -0.004 | 0.135 |
| BTN2A2 | 0.417 | 0.422 | -0.005 | 0.032 |
| WDR73 | 0.006 | 0.015 | -0.010 | 0.033 |
| LRRC37A2 | 0.124 | 0.145 | -0.021 | 0.007 |
| PPP2R3C | 0.269 | 0.304 | -0.035 | 0.091 |
| SRA1 | 0.458 | 0.497 | -0.038 | 0.049 |
| ERCC8 | 0.014 | 0.054 | -0.040 | 0.185 |
| MAPK3 | 0.304 | 0.353 | -0.050 | 0.272 |

Table S1: Data for all genes accompanying Figure 5B

### 9 EMBER and regression results for all genes

Figure S4: Shown are reported TWAS/GTEx v7 effects versus effects inferred from CommonMind gene expression data for all 36 genes with TWAS p-value $< 10^{-5}$, arrayed according to SNP position. For each gene, the top row displays TWAS/FUSION reported BLUP effects on GTEx v7 data; the middle row displays BLUP effects (blue) and when available, TWAS/FUSION reported BLUP effects on CMC data (black); and the bottom row displays results of EMBER (green) and linear regression (black), both using variable reduction to resolve high LD across SNP predictors.

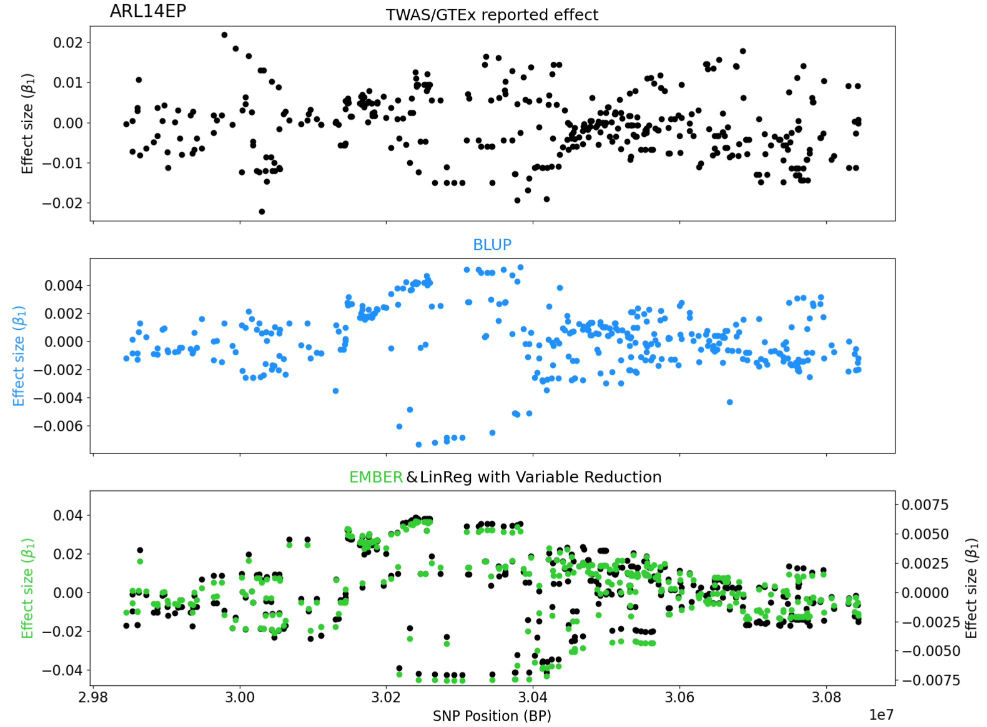

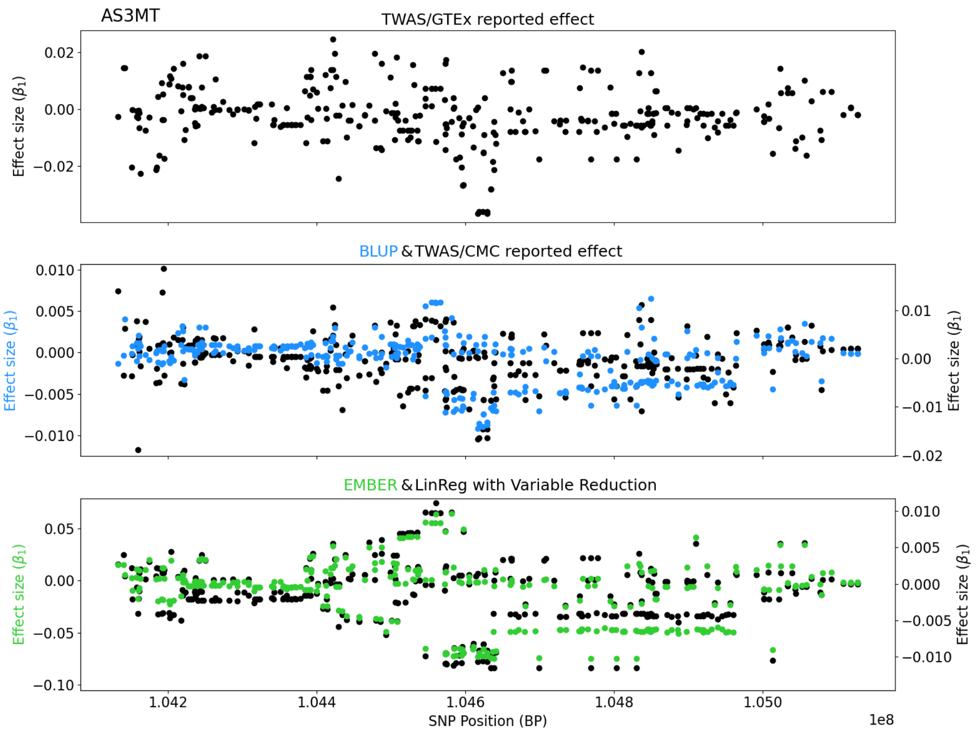
AS3MT
TWAS/GTEx reported effect
Effect size ($\beta_1$)
BLUP & TWAS/CMC reported effect
EMBER & LinReg with Variable Reduction
SNP Position (BP)
1e8

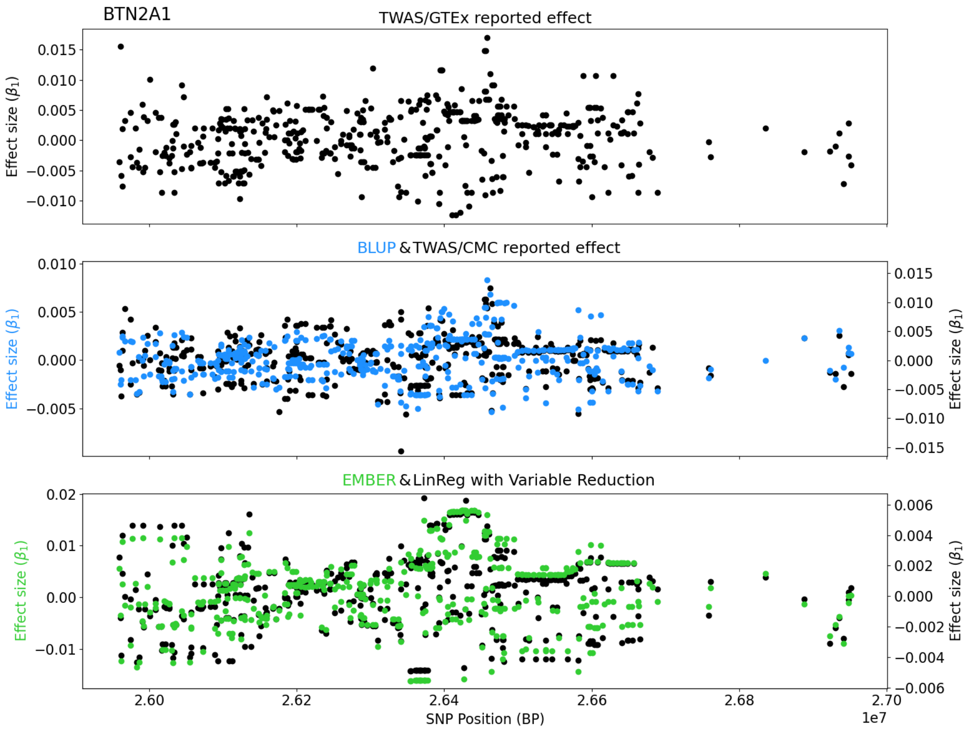
BTN2A1
TWAS/GTEx reported effect
Effect size ($\beta_1$)
BLUP & TWAS/CMC reported effect
EMBER & LinReg with Variable Reduction
SNP Position (BP)
1e7

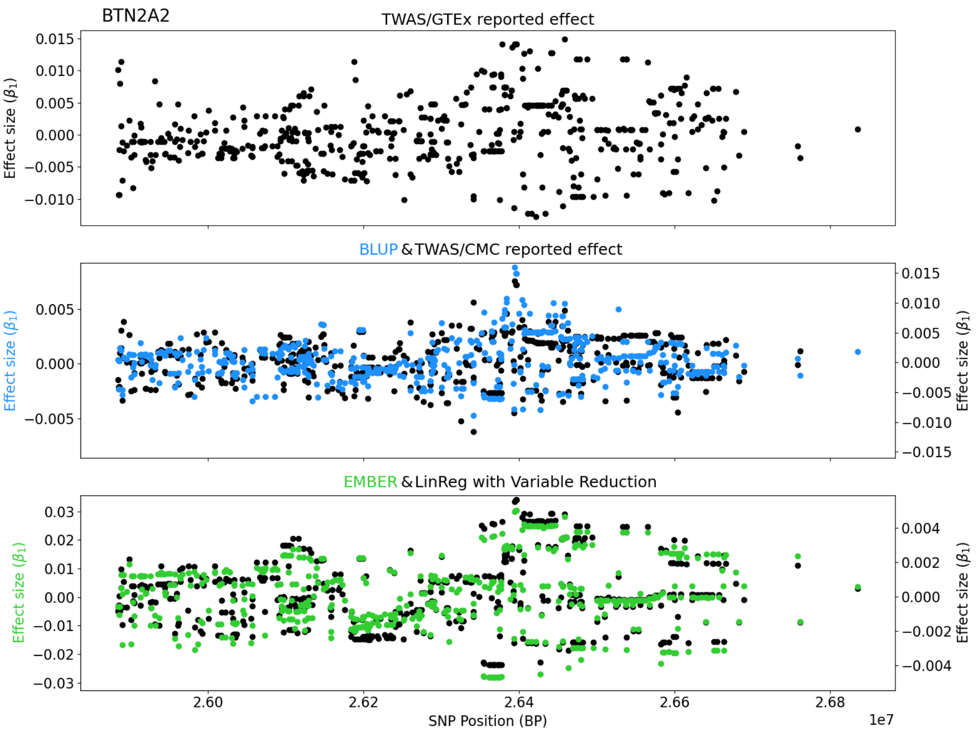
BTN2A2
TWAS/GTEx reported effect
BLUP & TWAS/CMC reported effect
EMBER & LinReg with Variable Reduction
Effect size ($\beta_1$)
SNP Position (BP)
1e7
2.60
2.62
2.64
2.66
2.68

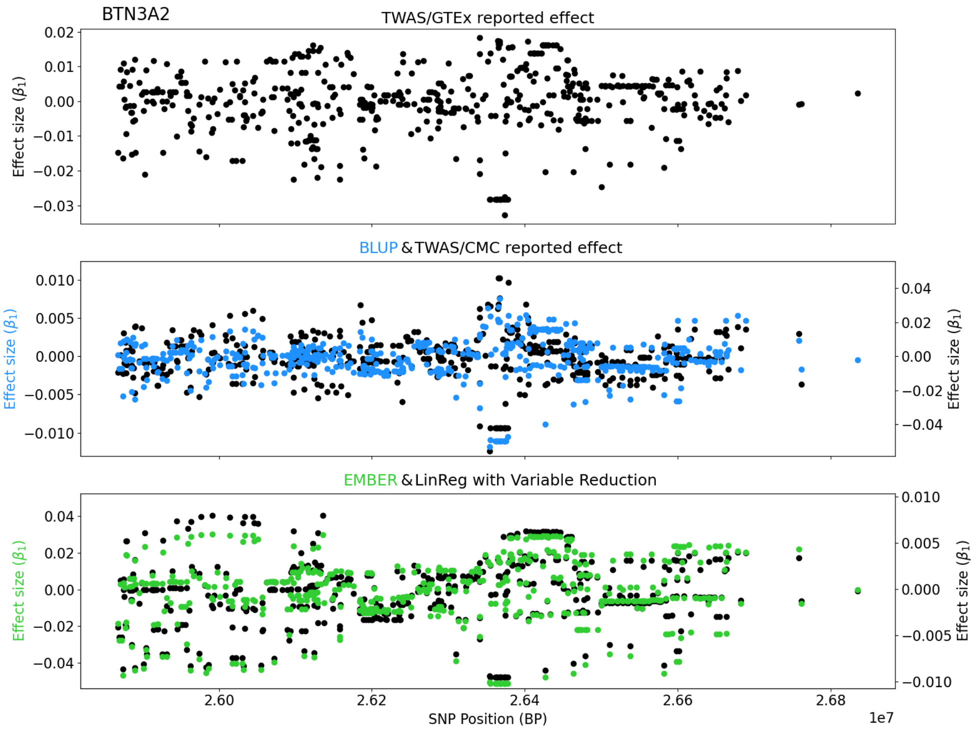
BTN3A2
TWAS/GTEx reported effect
BLUP & TWAS/CMC reported effect
EMBER & LinReg with Variable Reduction
Effect size ($\beta_1$)
SNP Position (BP)
1e7
2.60
2.62
2.64
2.66
2.68

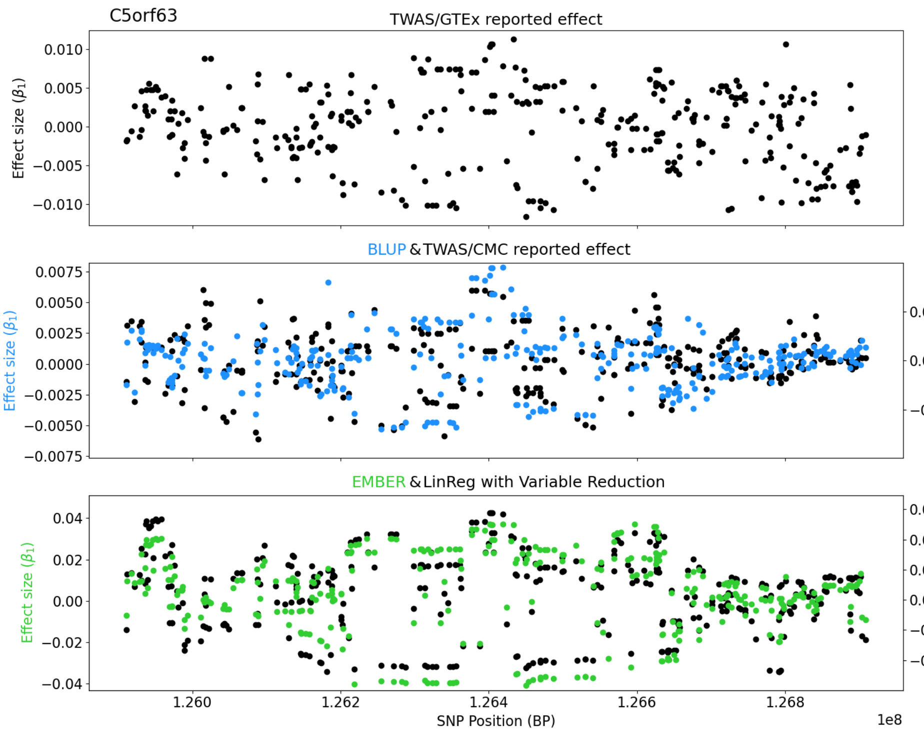
C5orf63
TWAS/GTEx reported effect
Effect size (β1)
BLUP & TWAS/CMC reported effect
EMBER & LinReg with Variable Reduction
SNP Position (BP)
1e8

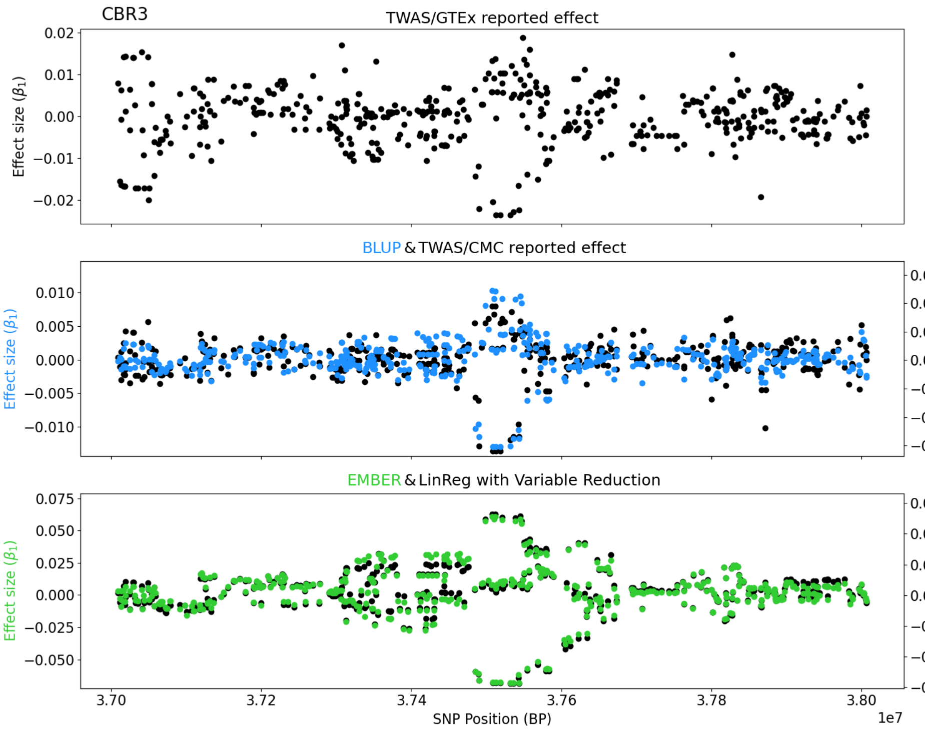
CBR3
TWAS/GTEx reported effect
Effect size (β1)
BLUP & TWAS/CMC reported effect
EMBER & LinReg with Variable Reduction
SNP Position (BP)
1e7

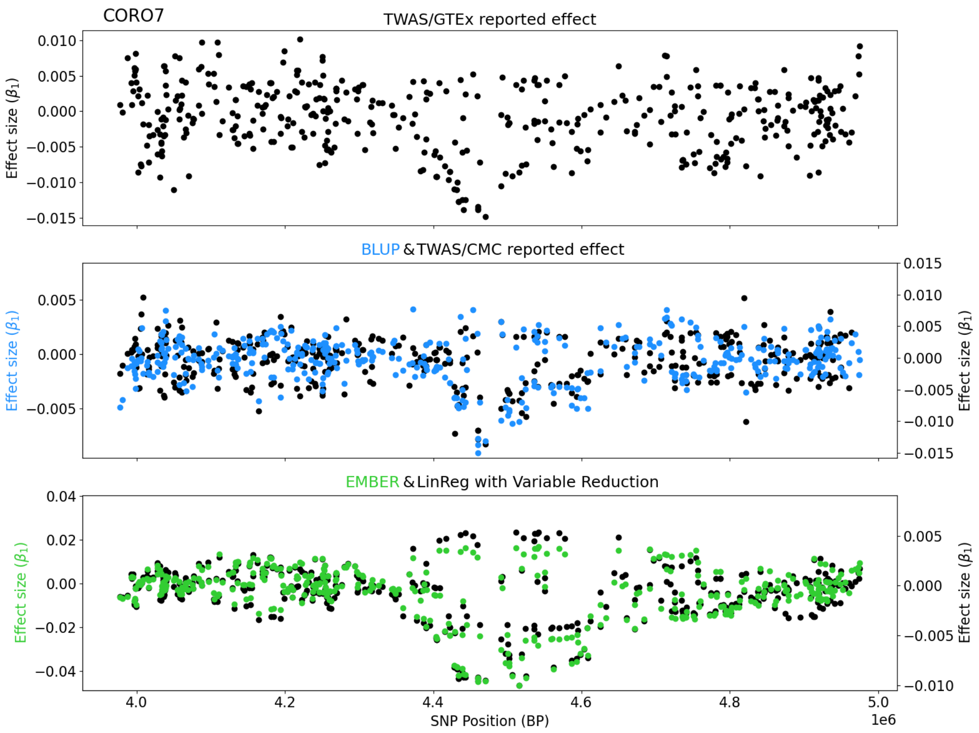
CORO7
TWAS/GTEx reported effect
Effect size ($\beta_1$)
BLUP & TWAS/CMC reported effect
Effect size ($\beta_1$)
Effect size ($\beta_1$)
EMBER & LinReg with Variable Reduction
Effect size ($\beta_1$)
Effect size ($\beta_1$)
SNP Position (BP)
1e6

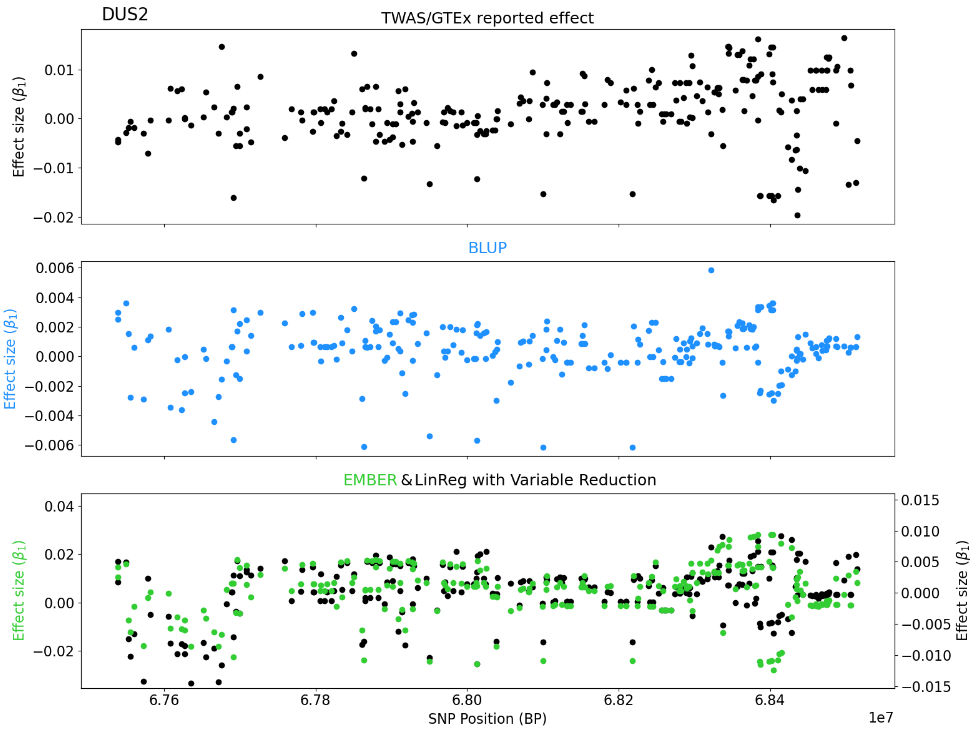
DUS2
TWAS/GTEx reported effect
Effect size ($\beta_1$)
BLUP
Effect size ($\beta_1$)
EMBER & LinReg with Variable Reduction
Effect size ($\beta_1$)
Effect size ($\beta_1$)
SNP Position (BP)
1e7

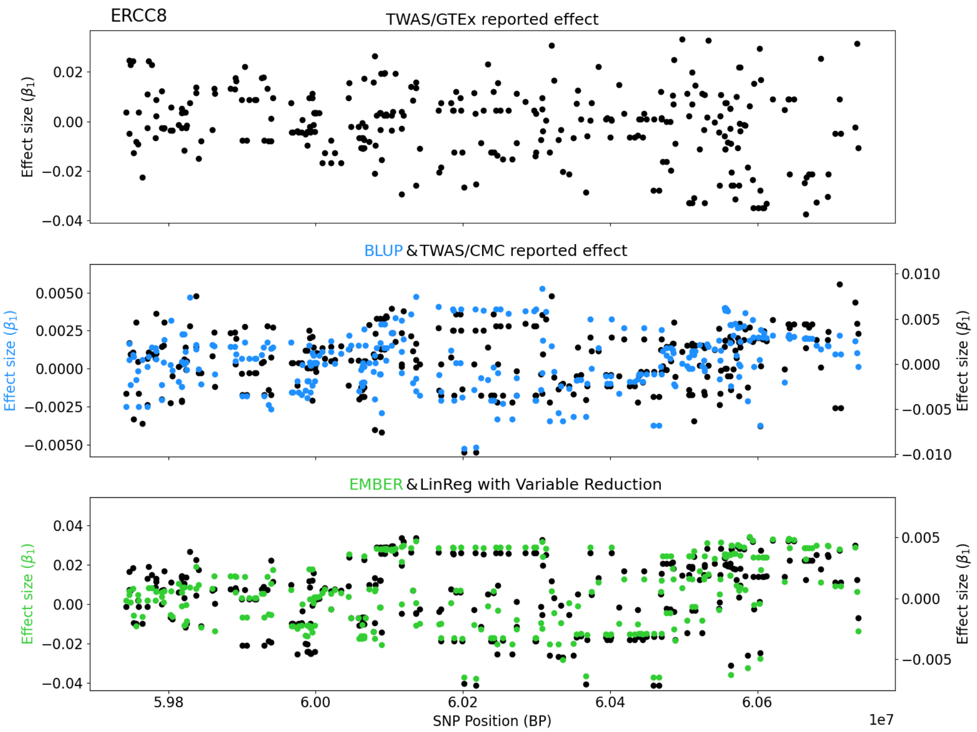
ERCC8
TWAS/GTEx reported effect
Effect size ($\beta_1$)
BLUP & TWAS/CMC reported effect
EMBER & LinReg with Variable Reduction
SNP Position (BP)
1e7

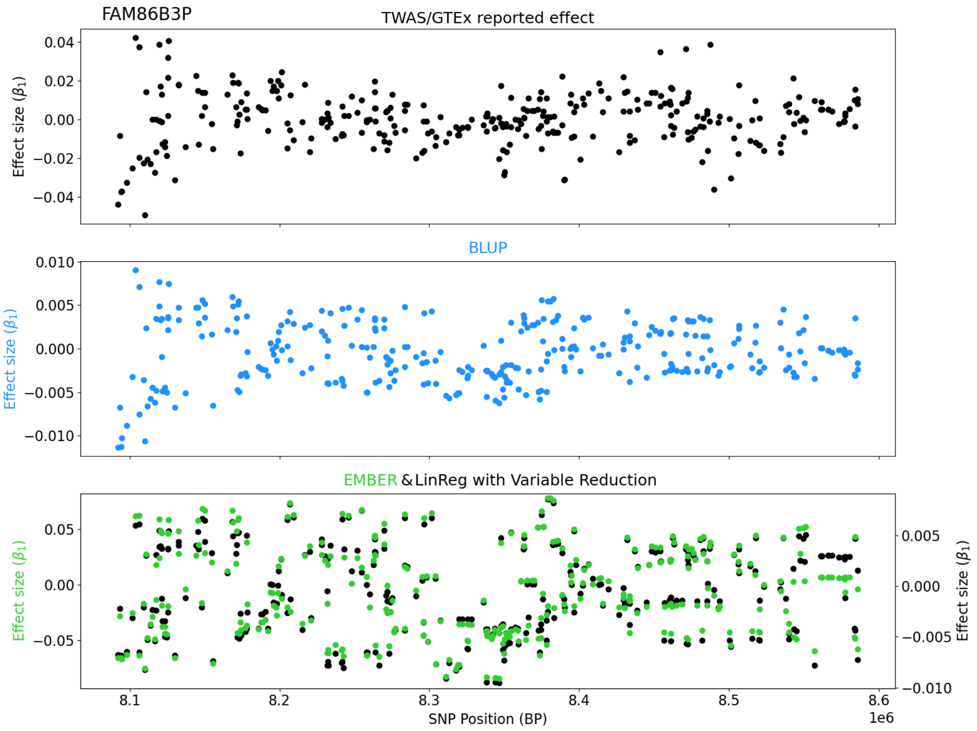
FAM86B3P
TWAS/GTEx reported effect
Effect size ($\beta_1$)
BLUP
EMBER & LinReg with Variable Reduction
SNP Position (BP)
1e6

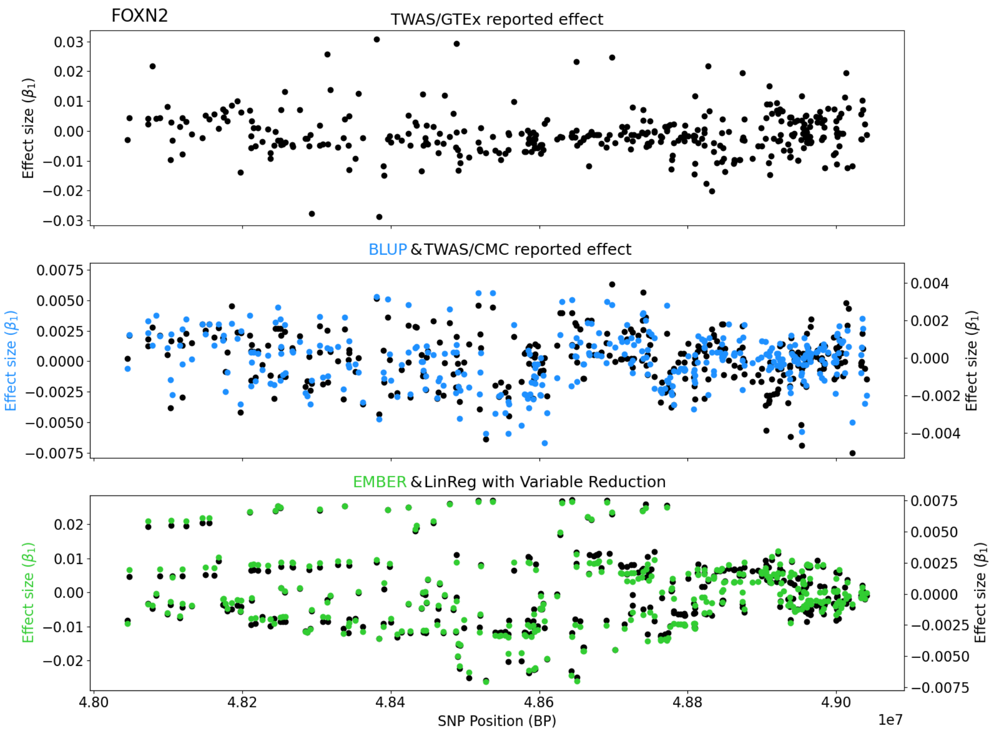
FOXN2
TWAS/GTEx reported effect
Effect size ($\beta_1$)
BLUP & TWAS/CMC reported effect
EMBER & LinReg with Variable Reduction
SNP Position (BP)
1e7

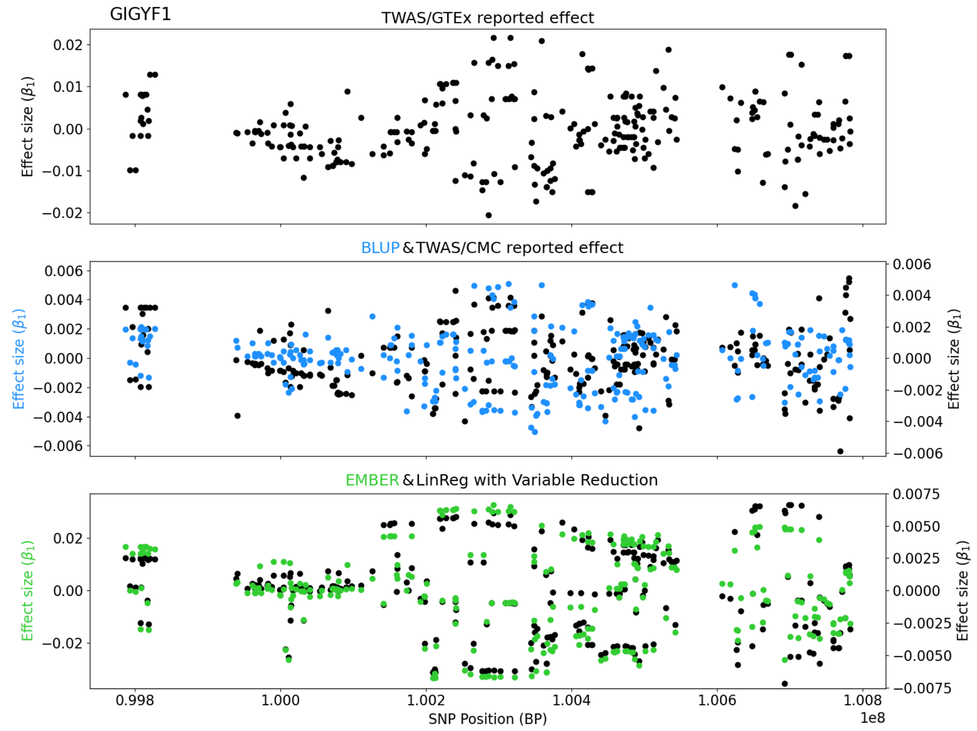
GIGYF1
TWAS/GTEx reported effect
Effect size ($\beta_1$)
BLUP & TWAS/CMC reported effect
EMBER & LinReg with Variable Reduction
SNP Position (BP)
1e8

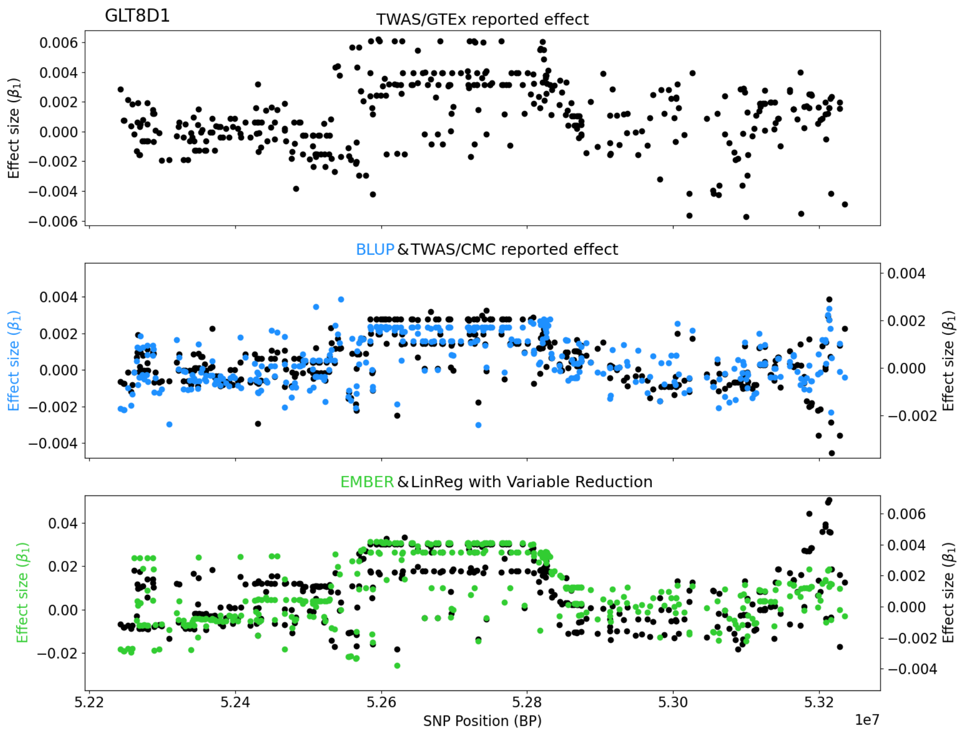
GLT8D1
TWAS/GTEx reported effect
Effect size ($\beta_1$)
BLUP & TWAS/CMC reported effect
Effect size ($\beta_1$)
Effect size ($\beta_1$)
EMBER & LinReg with Variable Reduction
Effect size ($\beta_1$)
Effect size ($\beta_1$)
SNP Position (BP)
1e7

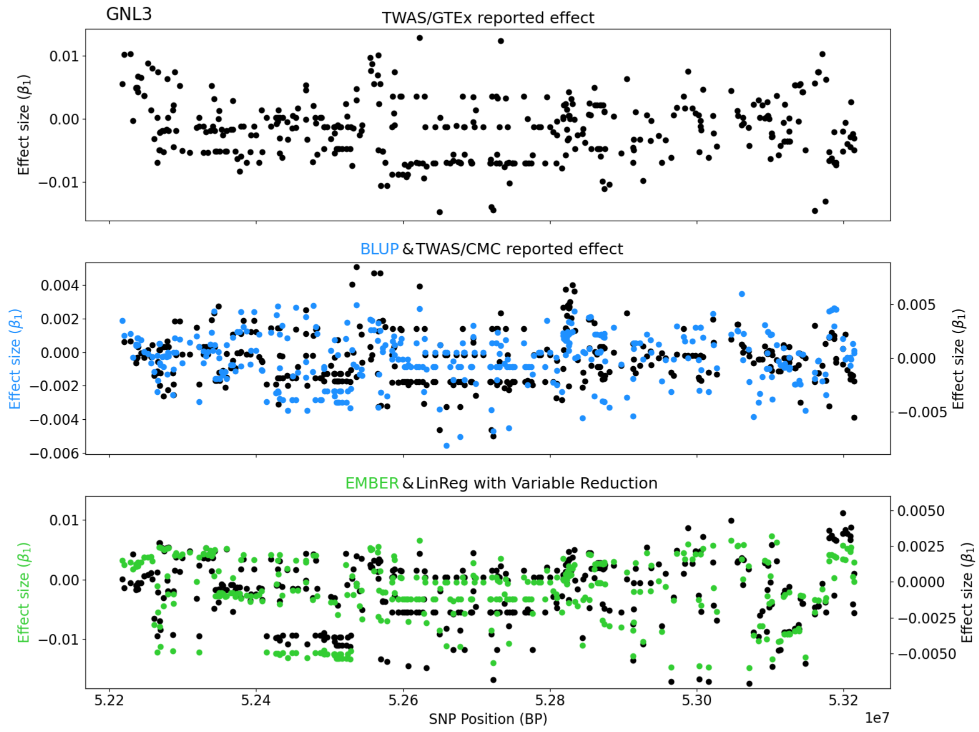
GNL3
TWAS/GTEx reported effect
Effect size ($\beta_1$)
BLUP & TWAS/CMC reported effect
Effect size ($\beta_1$)
Effect size ($\beta_1$)
EMBER & LinReg with Variable Reduction
Effect size ($\beta_1$)
Effect size ($\beta_1$)
SNP Position (BP)
1e7

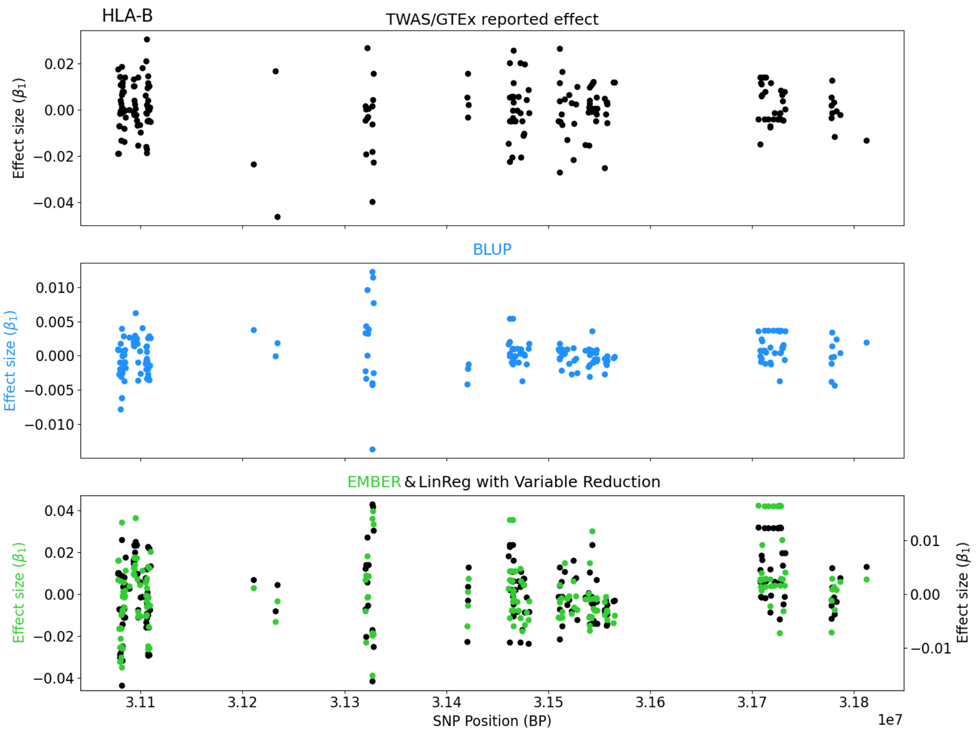
HLA-B
TWAS/GTEx reported effect
BLUP
EMBER & LinReg with Variable Reduction
Effect size ($\beta_1$)
SNP Position (BP)
1e7
3.11
3.12
3.13
3.14
3.15
3.16
3.17
3.18

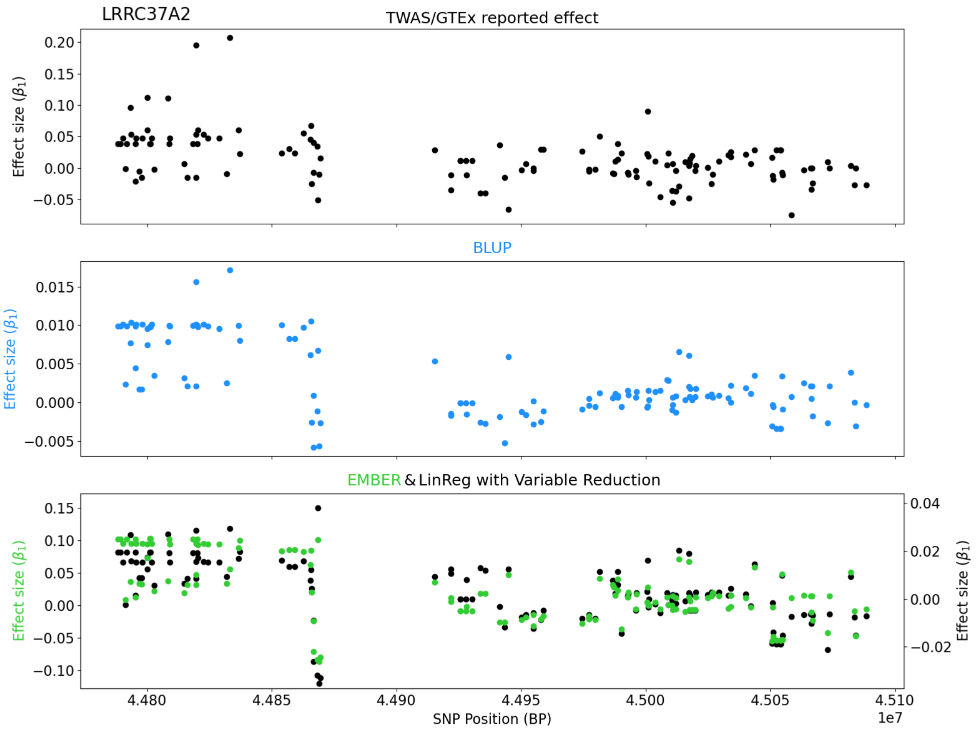
LRRC37A2
TWAS/GTEx reported effect
BLUP
EMBER & LinReg with Variable Reduction
Effect size ($\beta_1$)
SNP Position (BP)
1e7
4.480
4.485
4.490
4.495
4.500
4.505
4.510

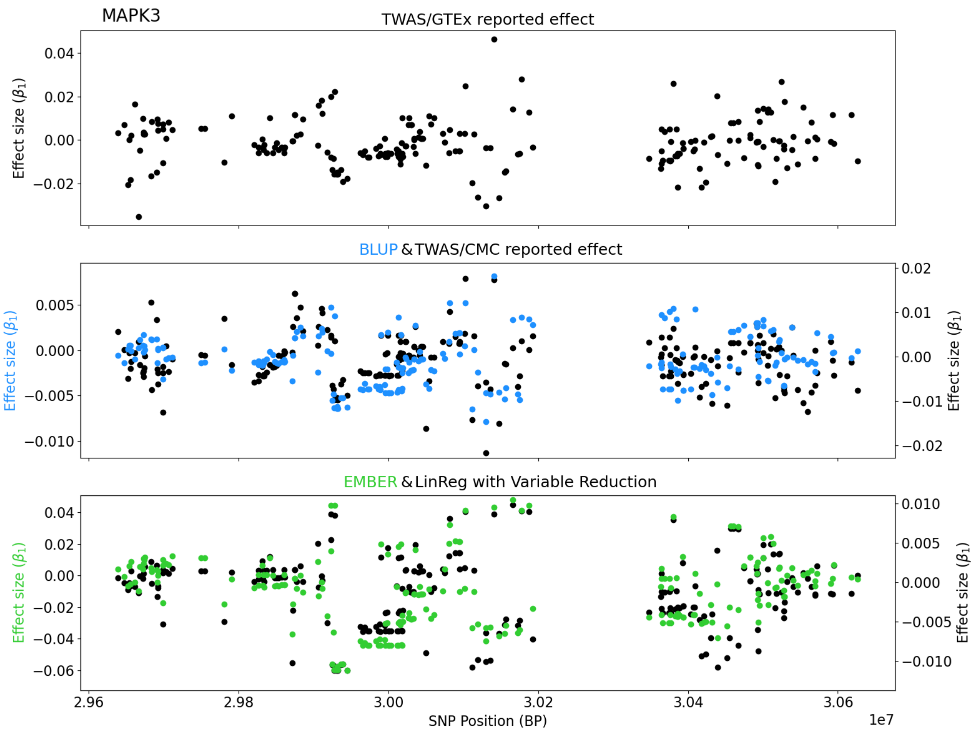
MAPK3
TWAS/GTEx reported effect
Effect size ($\beta_1$)
BLUP & TWAS/CMC reported effect
Effect size ($\beta_1$)
EMBER & LinReg with Variable Reduction
Effect size ($\beta_1$)
SNP Position (BP)
1e7

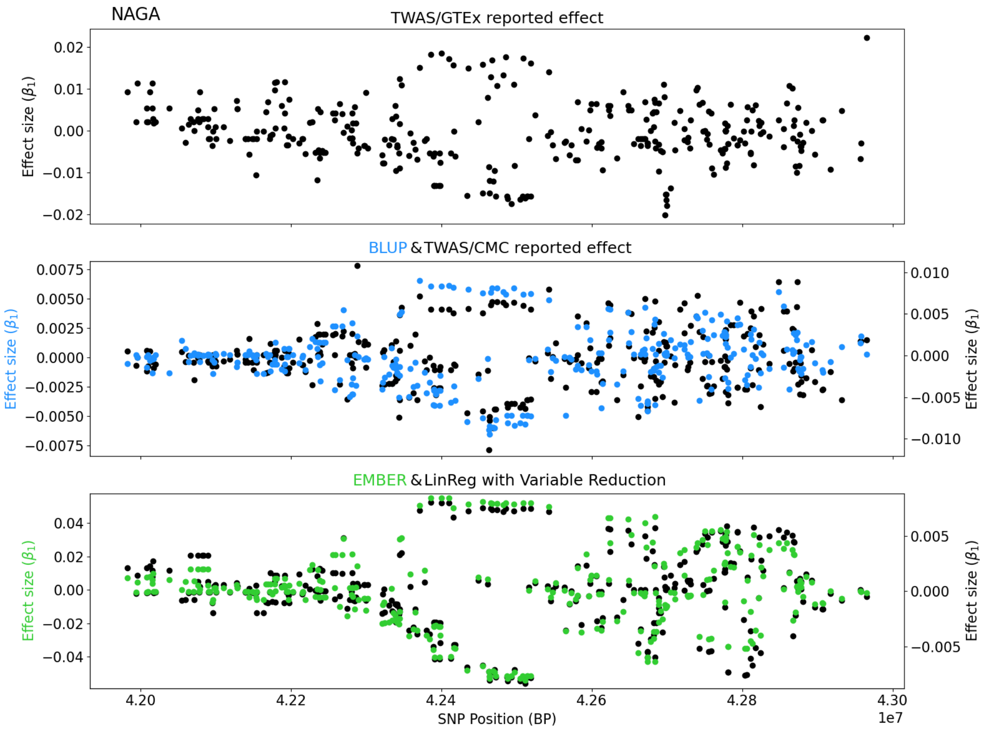
NAGA
TWAS/GTEx reported effect
Effect size ($\beta_1$)
BLUP & TWAS/CMC reported effect
Effect size ($\beta_1$)
EMBER & LinReg with Variable Reduction
Effect size ($\beta_1$)
SNP Position (BP)
1e7

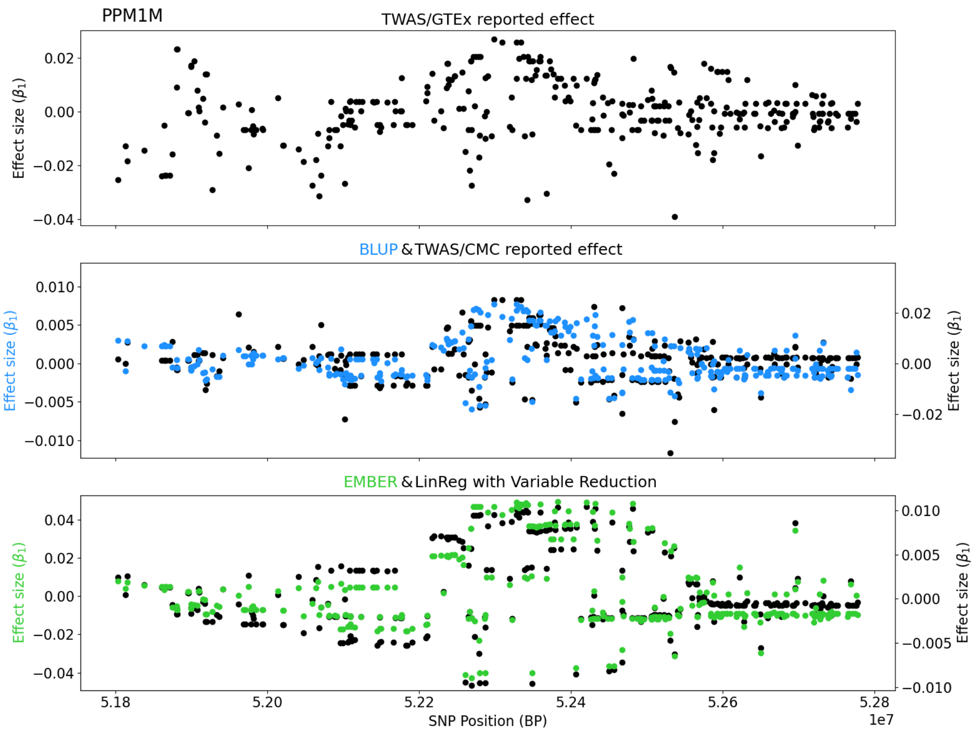
PPM1M
TWAS/GTEx reported effect
Effect size ($\beta_1$)
BLUP & TWAS/CMC reported effect
Effect size ($\beta_1$)
Effect size ($\beta_1$)
EMBER & LinReg with Variable Reduction
Effect size ($\beta_1$)
Effect size ($\beta_1$)
SNP Position (BP)
1e7

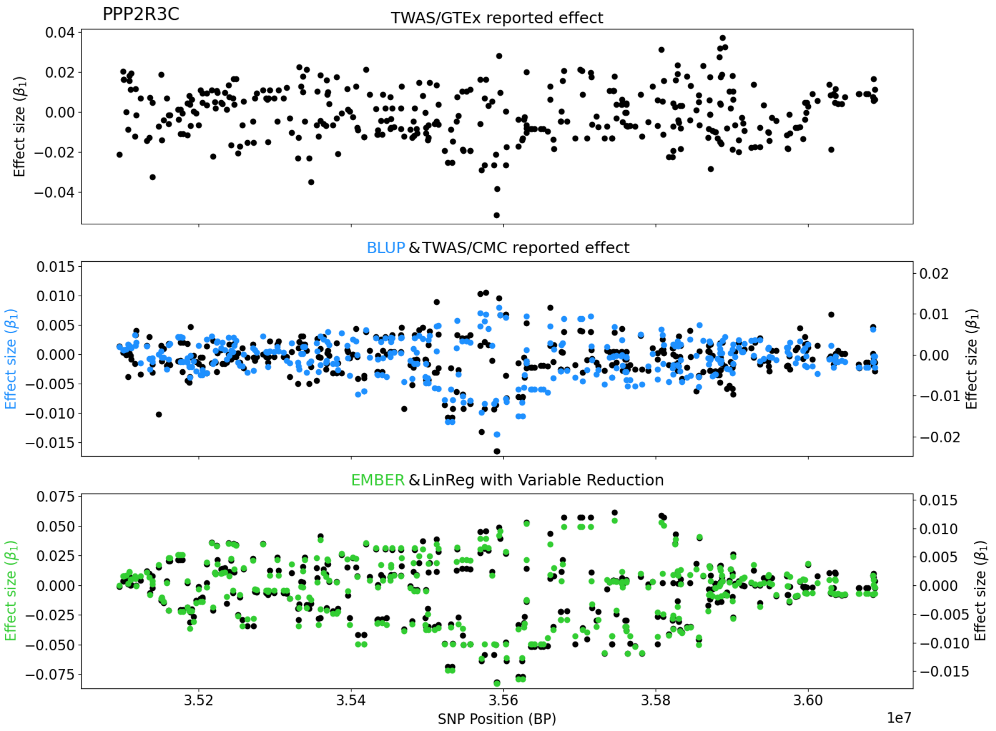
PPP2R3C
TWAS/GTEx reported effect
Effect size ($\beta_1$)
BLUP & TWAS/CMC reported effect
Effect size ($\beta_1$)
Effect size ($\beta_1$)
EMBER & LinReg with Variable Reduction
Effect size ($\beta_1$)
Effect size ($\beta_1$)
SNP Position (BP)
1e7

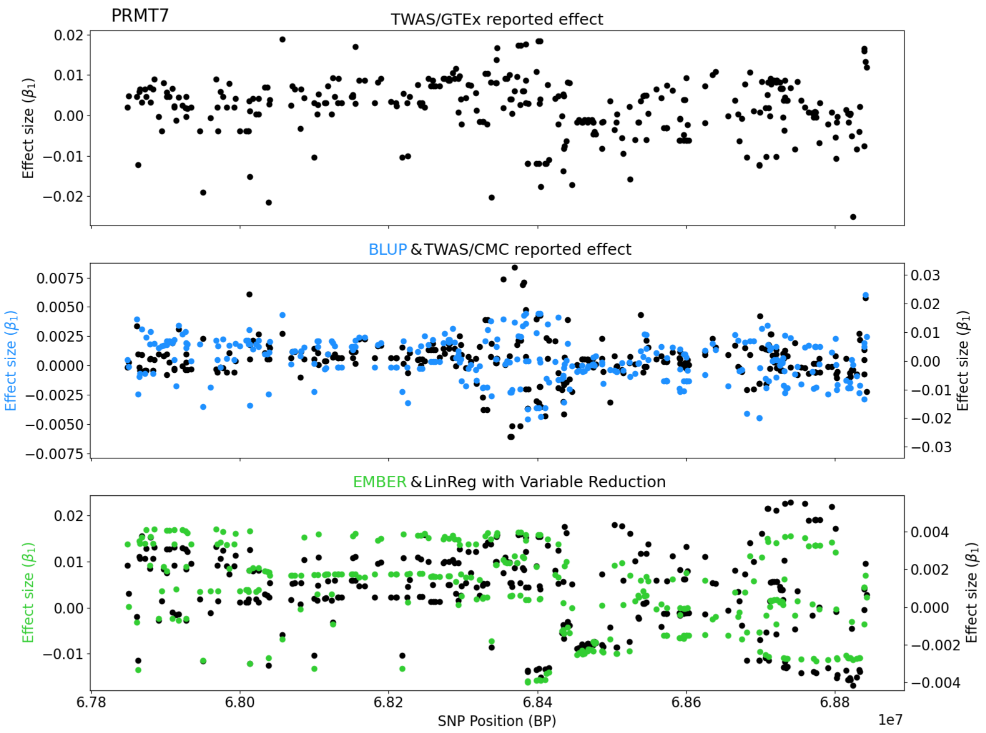
PRMT7
TWAS/GTEx reported effect
Effect size ($\beta_1$)
BLUP & TWAS/CMC reported effect
EMBER & LinReg with Variable Reduction
SNP Position (BP)
1e7

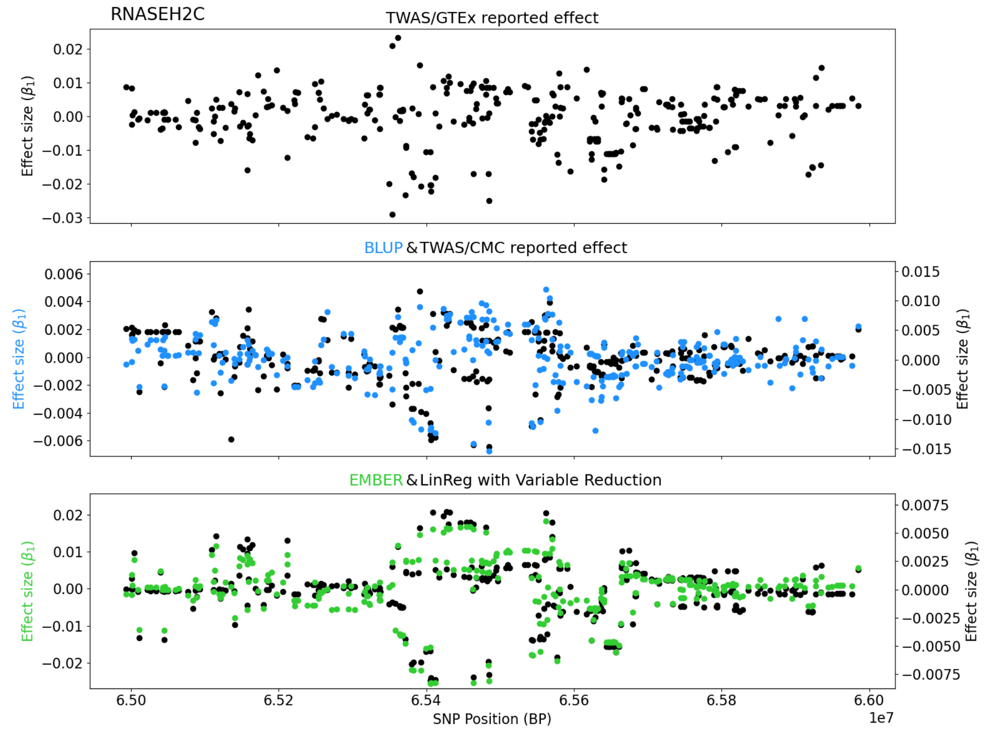
RNASEH2C
TWAS/GTEx reported effect
Effect size ($\beta_1$)
BLUP & TWAS/CMC reported effect
EMBER & LinReg with Variable Reduction
SNP Position (BP)
1e7

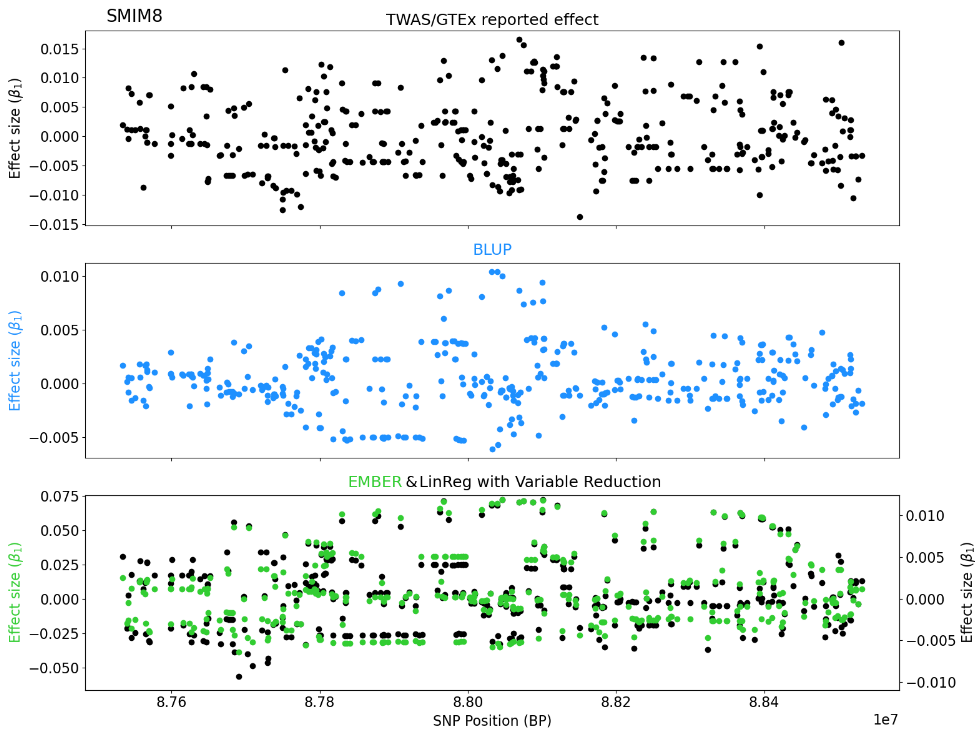
SMIM8
TWAS/GTEx reported effect
Effect size ($\beta_1$)
0.015
0.010
0.005
0.000
−0.005
−0.010
−0.015
BLUP
Effect size ($\beta_1$)
0.010
0.005
0.000
−0.005
EMBER & LinReg with Variable Reduction
Effect size ($\beta_1$)
0.075
0.050
0.025
0.000
−0.025
−0.050
Effect size ($\beta_1$)
0.010
0.005
0.000
−0.005
−0.010
8.76
8.78
8.80
8.82
8.84
1e7
SNP Position (BP)

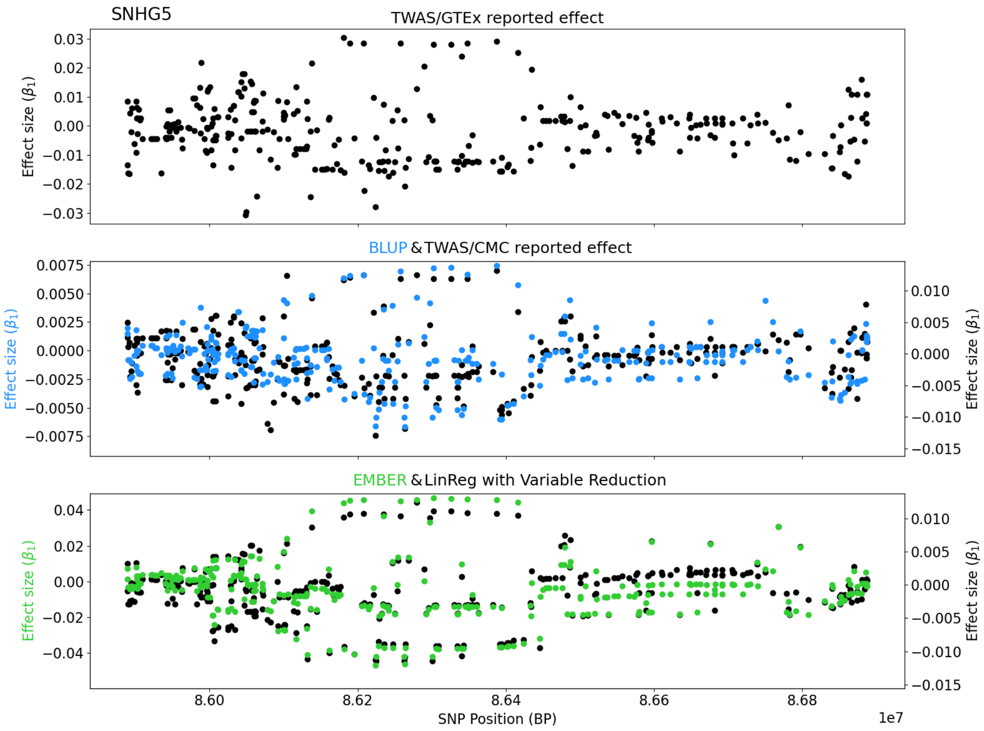
SNHG5
TWAS/GTEx reported effect
Effect size ($\beta_1$)
0.03
0.02
0.01
0.00
−0.01
−0.02
−0.03
BLUP & TWAS/CMC reported effect
Effect size ($\beta_1$)
0.0075
0.0050
0.0025
0.0000
−0.0025
−0.0050
−0.0075
Effect size ($\beta_1$)
0.010
0.005
0.000
−0.005
−0.010
−0.015
EMBER & LinReg with Variable Reduction
Effect size ($\beta_1$)
0.04
0.02
0.00
−0.02
−0.04
Effect size ($\beta_1$)
0.010
0.005
0.000
−0.005
−0.010
−0.015
8.60
8.62
8.64
8.66
8.68
1e7
SNP Position (BP)

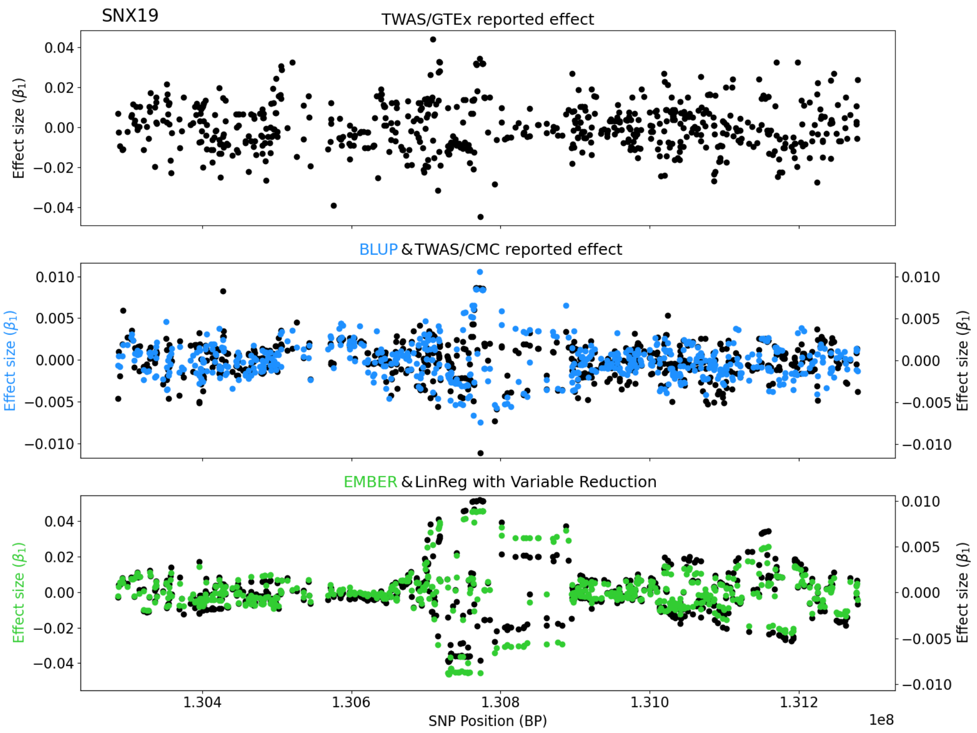
SNX19
TWAS/GTEx reported effect
Effect size ($\beta_1$)
BLUP & TWAS/CMC reported effect
Effect size ($\beta_1$)
Effect size ($\beta_1$)
EMBER & LinReg with Variable Reduction
Effect size ($\beta_1$)
Effect size ($\beta_1$)
SNP Position (BP)
1e8

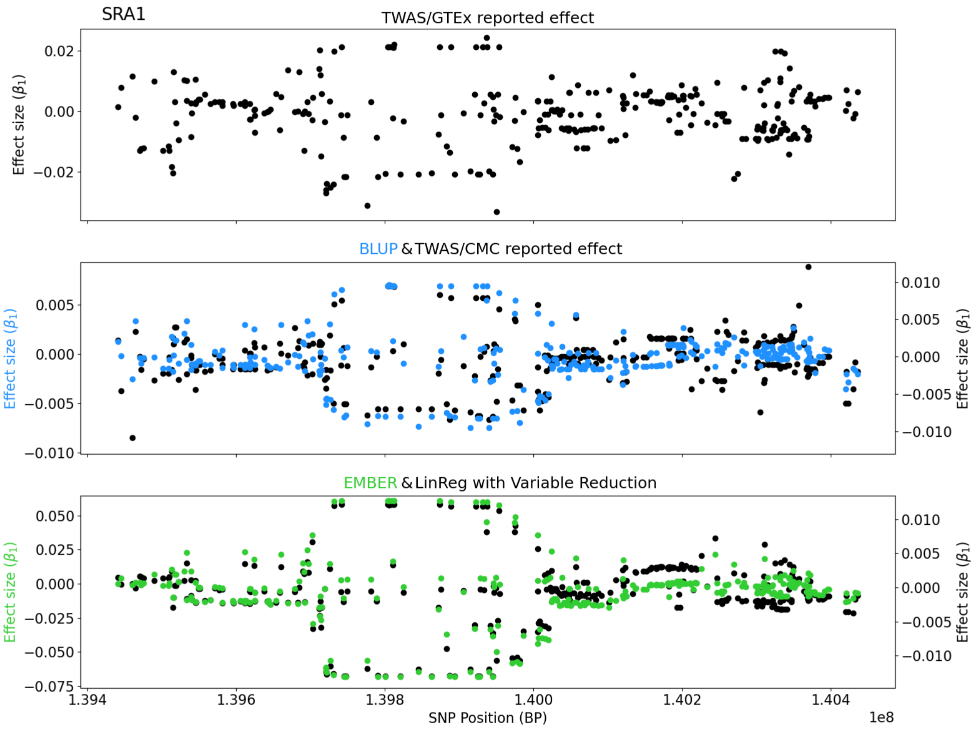
SRA1
TWAS/GTEx reported effect
Effect size ($\beta_1$)
BLUP & TWAS/CMC reported effect
Effect size ($\beta_1$)
Effect size ($\beta_1$)
EMBER & LinReg with Variable Reduction
Effect size ($\beta_1$)
Effect size ($\beta_1$)
SNP Position (BP)
1e8

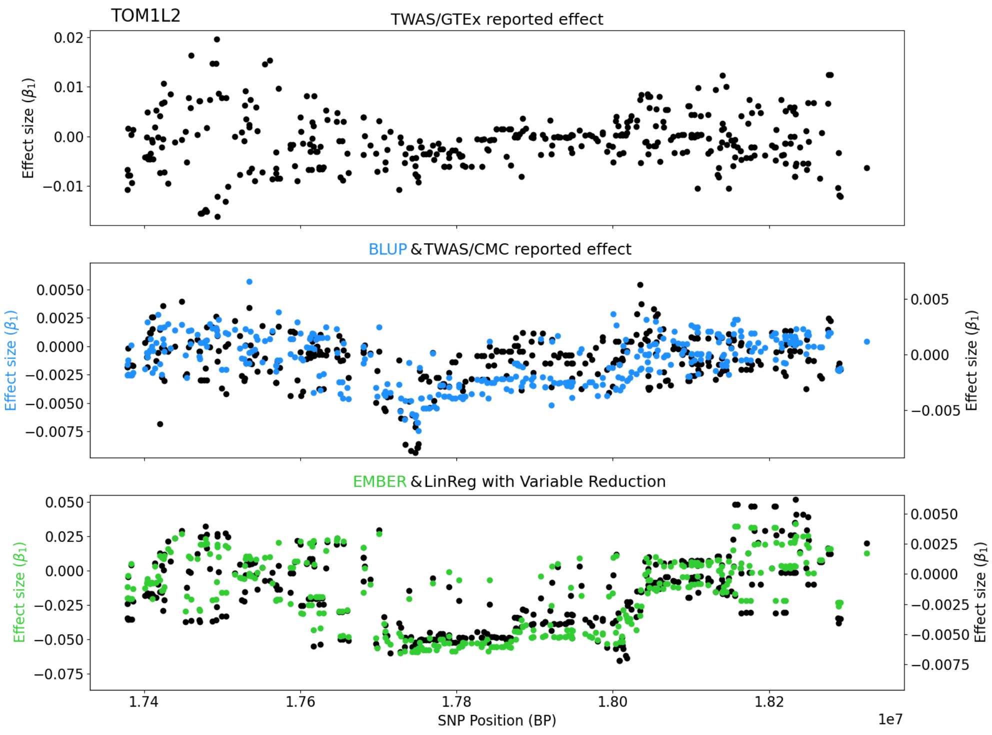
TOM1L2
TWAS/GTEx reported effect
Effect size ($\beta_1$)
0.02
0.01
0.00
−0.01
BLUP & TWAS/CMC reported effect
0.0050
0.0025
0.0000
−0.0025
−0.0050
−0.0075
0.005
0.000
−0.005
EMBER & LinReg with Variable Reduction
0.050
0.025
0.000
−0.025
−0.050
−0.075
0.0050
0.0025
0.0000
−0.0025
−0.0050
−0.0075
1.74
1.76
1.78
1.80
1.82
SNP Position (BP)
1e7

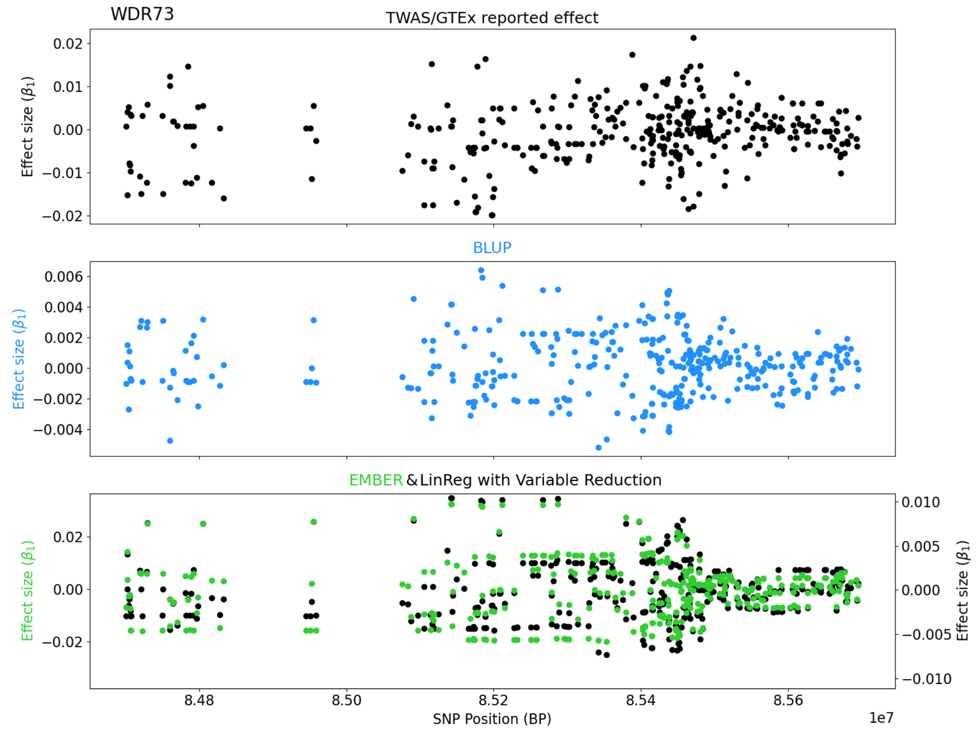
WDR73
TWAS/GTEx reported effect
Effect size ($\beta_1$)
0.02
0.01
0.00
−0.01
−0.02
BLUP
0.006
0.004
0.002
0.000
−0.002
−0.004
EMBER & LinReg with Variable Reduction
0.02
0.00
−0.02
0.010
0.005
0.000
−0.005
−0.010
8.48
8.50
8.52
8.54
8.56
SNP Position (BP)
1e7

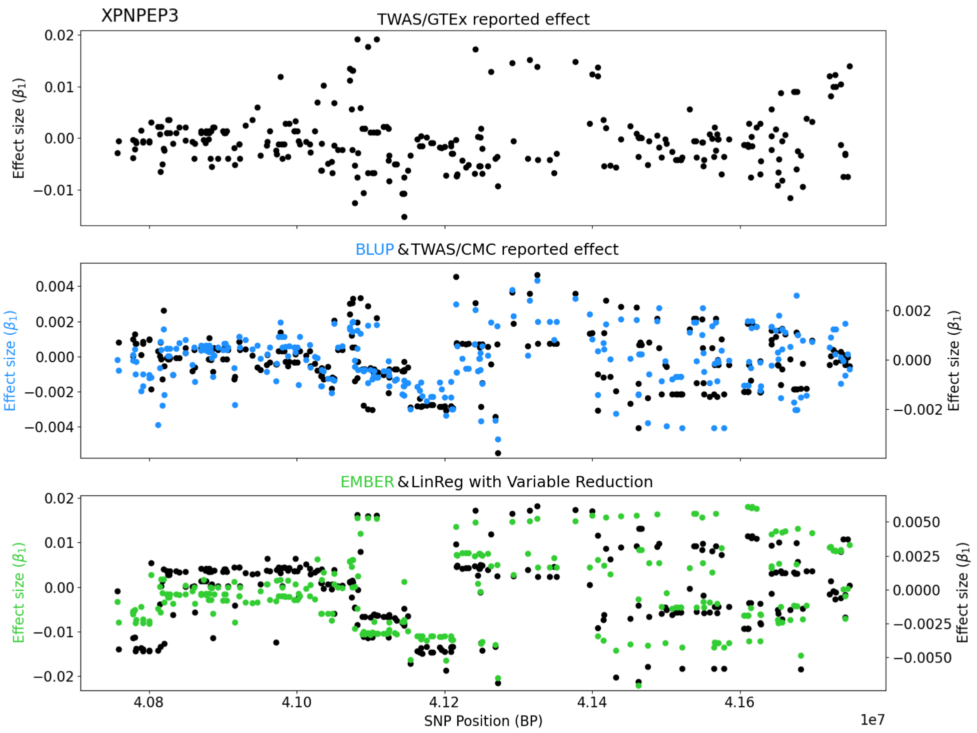
XPNPEP3
TWAS/GTEx reported effect
BLUP & TWAS/CMC reported effect
EMBER & LinReg with Variable Reduction
Effect size ($\beta_1$)
SNP Position (BP)
1e7
4.08
4.10
4.12
4.14
4.16

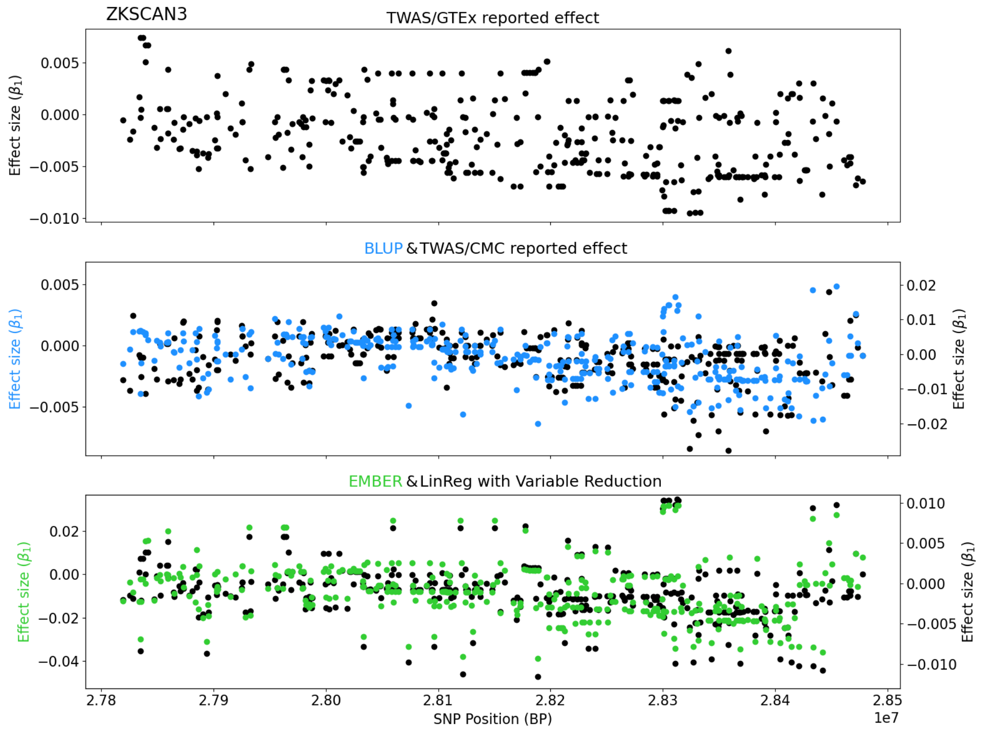
ZKSCAN3
TWAS/GTEx reported effect
BLUP & TWAS/CMC reported effect
EMBER & LinReg with Variable Reduction
Effect size ($\beta_1$)
SNP Position (BP)
1e7
2.78
2.79
2.80
2.81
2.82
2.83
2.84
2.85

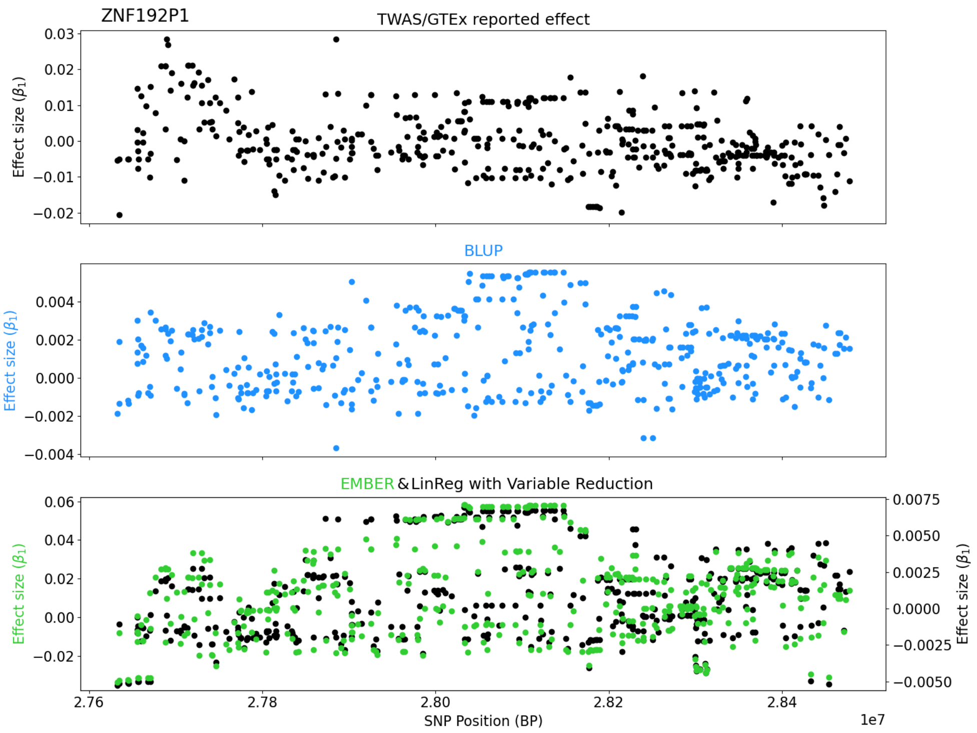
ZNF192P1
TWAS/GTEx reported effect
Effect size ($\beta_1$)
BLUP
Effect size ($\beta_1$)
EMBER & LinReg with Variable Reduction
Effect size ($\beta_1$)
Effect size ($\beta_1$)
SNP Position (BP)
1e7

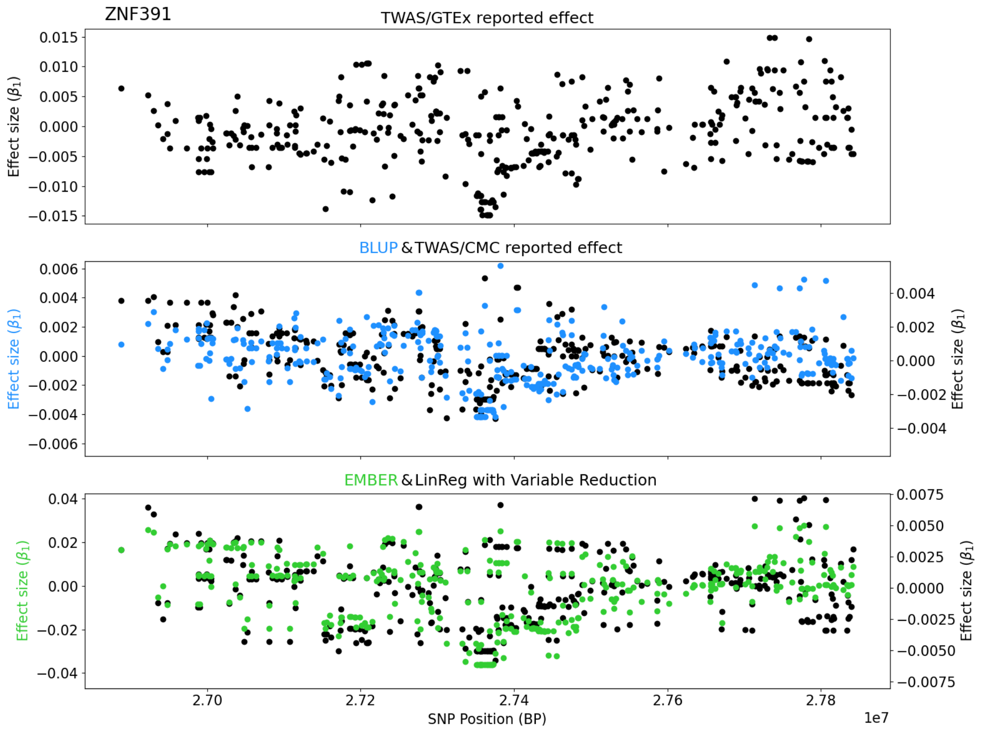
ZNF391
TWAS/GTEx reported effect
Effect size ($\beta_1$)
BLUP & TWAS/CMC reported effect
Effect size ($\beta_1$)
Effect size ($\beta_1$)
EMBER & LinReg with Variable Reduction
Effect size ($\beta_1$)
Effect size ($\beta_1$)
SNP Position (BP)
1e7

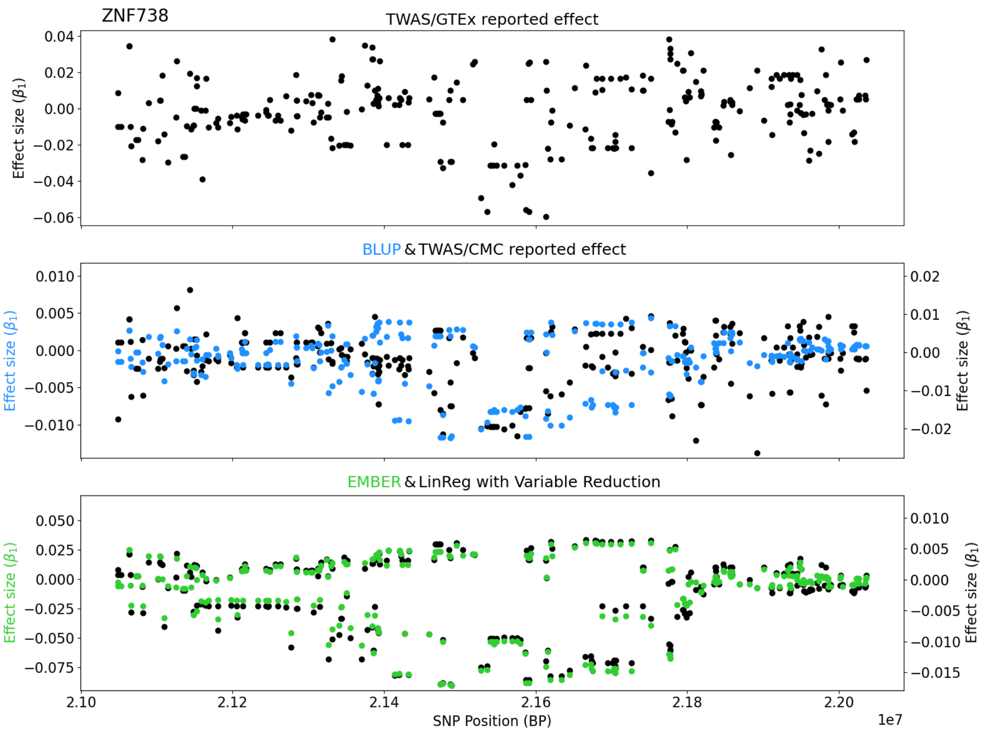

ZNF738
TWAS/GTEx reported effect
Effect size ($\beta_1$)
BLUP & TWAS/CMC reported effect
EMBER & LinReg with Variable Reduction
SNP Position (BP)
1e7


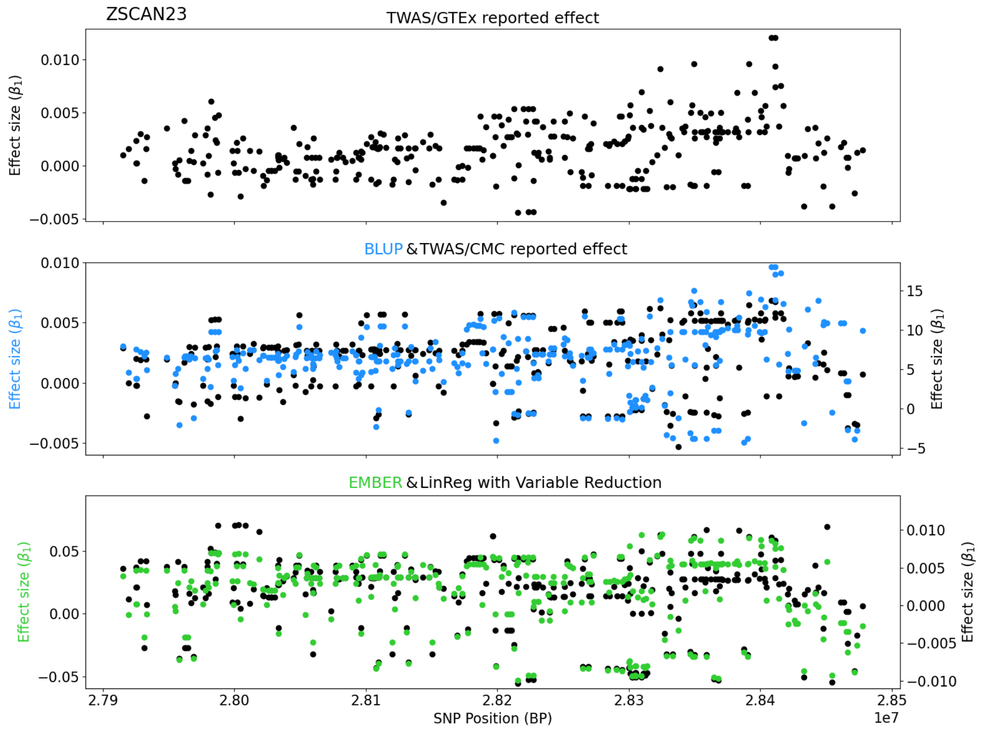

ZSCAN23
TWAS/GTEx reported effect
Effect size ($\beta_1$)
BLUP & TWAS/CMC reported effect
EMBER & LinReg with Variable Reduction
SNP Position (BP)
1e7

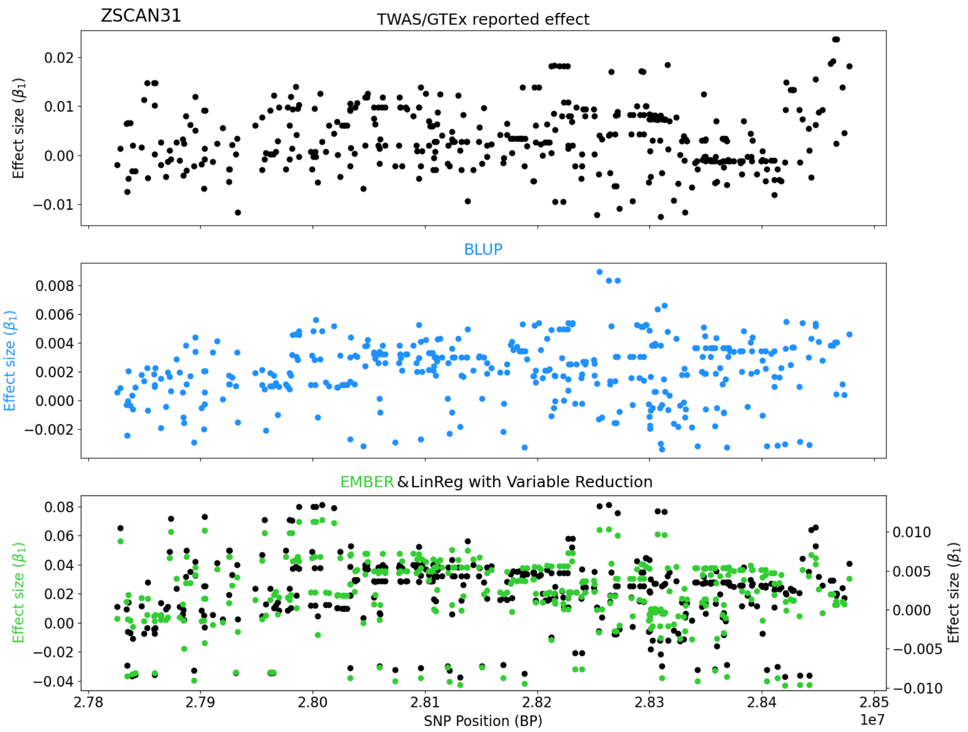
ZSCAN31
TWAS/GTEx reported effect
Effect size ($\beta_1$)
0.02
0.01
0.00
−0.01
BLUP
Effect size ($\beta_1$)
0.008
0.006
0.004
0.002
0.000
−0.002
EMBER & LinReg with Variable Reduction
Effect size ($\beta_1$)
0.08
0.06
0.04
0.02
0.00
−0.02
−0.04
Effect size ($\beta_1$)
0.010
0.005
0.000
−0.005
−0.010
2.78
2.79
2.80
2.81
2.82
2.83
2.84
2.85
1e7
SNP Position (BP)
